# Engineering reduced activity in the oxygen-sensing *Arabidopsis thaliana* plant cysteine oxidase 4 enzyme results in improved flood resilience

**DOI:** 10.1101/2025.07.24.666444

**Authors:** Anna Dirr, Arianna Del Greco, Simon Howe, Leah S. Walker, Monica Perri, Vinay Shukla, Francesco Licausi, Emily Flashman

## Abstract

Plant cysteine oxidases (PCOs) are O_2_-sensing enzymes that play an important role in plant responses to low oxygen (hypoxia). PCO-catalysed dioxygenation of the N-terminal Cys of substrates, including Group VII Ethylene Response Factors (ERVIIs), targets them for degradation via the Cys/Arg N-degron pathway, however these substrates are stabilized in hypoxia due to reduced PCO activity. When plants are flooded, submergence-induced hypoxia results in ERFVII-mediated upregulation of hypoxia responsive genes that reconfigure plant metabolism and allow short-term resilience to the conditions. However, the increasing frequency and duration of flood events requires strategies to improve plant flood resilience, particularly amongst agronomic crops. One possibility is to prolong the stability of ERFVIIs by engineering the PCOs to catalyse their oxidation less efficiently. We report a structure-guided kinetic and biophysical investigation of *Arabidopsis thaliana* PCO4 that reveals residues important for substrate-binding and catalysis. We subsequently selected *At*PCO4 variants Y183F and C173A, with severe and mild impacts on *At*PCO4 activity, respectively, to complement Arabidopsis *pco1pco2pco4pco5* plants and investigate their impact on submergence resilience. Both variants appeared to be beneficial for survival and recovery after 2.5 and 3.5 days of dark submergence when compared to control plants, indicating that engineering PCOs can be used as a strategy to improve flood tolerance in plants.

## Introduction

Floods are becoming more regular due to extreme weather events. While some agronomical plants (such as rice and some wheat cultivars) can tolerate waterlogged roots and submerged aerial parts, many commercially relevant crop plants are less tolerant, including barley, maize and soybean.^1,2^ From 2008 to 2018 floods resulted in USD 21 billion loss, the largest reduction in agricultural profit due to natural disaster after droughts.^3^ Hence, there is an urgent need to find ways to improve crop flood tolerance for future food security.

When plants are submerged, they experience multiple stresses: accumulated heavy metals in the soil, entrapped gasses such as ethylene and CO_2_, accumulation of reactive oxygen species (ROS) and reduced O_2_ availability (hypoxia) within the plant tissues due to slower diffusion and impaired photosynthesis.^4,5^ Ethylene and hypoxia trigger survival strategies within a submerged plant. Accumulated ethylene induces the transcription of *Group VII Ethylene Response Factors* (*ERFVII*s).^6^ ERFVIIs are normally degraded in normoxia through the activity of oxygen-dependent plant cysteine oxidases (PCOs) and the Cys/Arg-N-degron pathway:^7–9^ PCOs catalyse the dioxygenation of N-terminal Cys (Nt-Cys) of ERFVIIs (as well as other substrates^10,11^) to cysteine sulfinic acid (Cys-SO_2_H).^12^ The dioxygenated Nt-Cys is recognised by arginyl transferases (ATE) which catalyse the addition of an Arg to the respective ERFVII N-terminus,^12^ labelling them for ubiquitination by the E3 ligase Proteolysis 6 (PRT6) and proteasomal degradation.^13,14^ In contrast, during hypoxia PCO activity is limited, reducing Nt-Cys oxidation and leading to ERFVII stabilisation.^7–9^ ERFVIIs can then induce the transcription of hypoxia response genes (HRGs), including *alcohol dehydrogenase 1* (*ADH1*), *pyruvate decarboxylase 1* (*PDC1*) and *sucrose synthase 4* (*SUS4*), that support a shift to anaerobic metabolism to maintain basal levels of ATP production and allow temporary survival of flooding.^7,8,15,16^ PCO activity is therefore a key element in plant submergence resilience as it regulates the stability of ERFVIIs in response to O_2_ levels.

ERFVII accumulation and function is important both during submergence and upon de-submergence.^5,17^ As well as being stabilised by hypoxia, ERFVIIs are also impacted by other signals: (i) reduced ATP levels trigger the release of the Arabidopsis ERFVII Related to AP2.12 (RAP2.12) from its membrane-anchor,^18^ (ii) increased Ca^2+^ levels promote Thr phosphorylation in RAP2.12, RAP2.2 and RAP2.3, stabilising these ERFVIIs in the nucelus^19^ and (iii) nitric oxide (NO) scavenging by plant haemoglobin 1 / phytoglobin 1 (HB1 / PGB1, one of the HRGs) has been linked to the stabilisation of ERFVIIs via the N-degron pathway, independent of PCOs.^6,20,21^ The activity and accumulation of ERFVIIs is also regulated by negative feedback-loops from pathways sensing O_2_ concentrations (via induction of *AtPCO1* and *AtPCO2*, themselves HRGs)^9,16^ and energy levels^22–24^. When a flood recedes, the plant is exposed to new stresses including elevated ROS and photoinhibition, dehydration and nutrient deficiency.^5,17,25–31^ Acclimation strategies include the scavenging of ROS and tight regulation of stomata.^25,27,32,33^ In rice, the latter is promoted by SUB1A, an ERFVII in rice.^27^ Despite an Nt-Cys, SUB1A is not a substrate of PCOs due to its hidden Nt-Cys,^34^ wherefore SUB1A is stable during de-submergence and can confer submergence tolerance.^27^ This has been exploited to create agronomical rice cultivars with higher submergence tolerance.^35,36^ ERFVIIs are therefore fundamental to survival and recovery from submergence. In Arabidopsis it has been shown that transient accumulation of ERFVIIs can be beneficial for submergence survival, with similar outcomes reported in barley and maize.^8,37–39^ These observations suggest that fine-tuning ERFVII levels across the duration of a flood event, from the onset of hypoxia and upon reoxygenation, could therefore be a valuable tool to improve the hypoxia and submergence tolerance of plants. We have previously proposed that engineering PCOs to limit their rate of activity and enable ERFVII stabilisation in an O_2_ dependent manner could be a promising way of achieving this.^40^

PCOs belong to the Fe(II)-dependent thiol dioxygenase family which can be distinguished from other oxygenases by their conserved His-triad that coordinates the iron cofactor in the active site (Figure S1).^15,41^ Although an exact mechanism is not certain, it is considered likely, based on studies in other thiol dioxygenases, that PCO enzymes coordinate substrates via the substrates’ Nt-Cys amino and thiol groups at the active site Fe(II), prior to O_2_ binding and activation, allowing Nt-Cys oxidation via a Fe(III)-superoxo or Fe(IV)-oxo intermediate.^42,43^ We have previously reported *At*PCO4 variants that resulted in ablated function, both biochemically and when the variant *At*PCO4 sequences were introduced into *pco1pco2pco4pco5* Arabidopsis plants (hereafter *4pco*).^15^ In this study, we report the use of a structure/function-guided approach to design and test a suite of *At*PCO4 variants with reduced, but not ablated biochemical function. Based on comparison of the *At*PCO4 structure with related enzymes human 2-ethanethiol dioxygenase (*Hs*ADO) and human cysteine dioxygenase (*Hs*CDO-1),^15,41,42,44,45^ we mutated active site residues predicted to reduce activity via altered iron coordination, substrate affinity or catalytic rate (Figure S1). Following biochemical and kinetic characterisation of a suite of variants, we selected *At*PCO4 mutations C173A and Y183F to complement the Arabidopsis *4pco* model. We found that, in comparison to wildtype controls, plants containing either of these *At*PCO4 variants demonstrated improved recovery after dark submergence, confirming that engineering PCO function is a viable route to improve submergence tolerance in plants.

## Results and Discussion

### Selection of *At*PCO4 isoform for variant generation

Before designing *At*PCO4 variants, we wanted to ensure we were using the correct *AtPCO4* WT sequence for the Arabidopsis Columbia-0 (Col-0) plants in our laboratory because *At*PCO4 can be expressed as two splice variants, *At*PCO4-1 and *At*PCO4-2. *At*PCO4-2 contains an additional Glu residue at position 135 (E135) that is located in a loop away from the active site (Figure S2A). Col-0 plants in our laboratory showed higher transcription levels of *AtPCO4-2* (around 70%) than *AtPCO4-1* (around 30%). Since previous *in vitro* data focussed on *At*PCO4-1,^15^ we examined whether there are differences in *At*PCO-1 and *At*PCO4-2 activities *in vitro*, using previously reported methodology.^46,47^ We did not observe any significant difference in the activity of both *At*PCO4 isoforms (Figure S2) allowing the comparison of *in vitro* data from *At*PCO4-2 with *At*PCO4-1. Since Col-0 plants in our laboratory showed higher transcription levels of *AtPCO4-2* than *AtPCO4-1*, *At*PCO4-2 was used in this study to investigate active site residues *in vitro* and in the *4pco* Arabidopsis model. Herein, *At*PCO4-2 will be referred to as *At*PCO4.

### Replacement of iron-coordinating residues ablates *At*PCO4 activity

All *At*PCOs, in common with other thiol dioxygenases, contain a conserved iron coordinating His-triad (Figure S1C).^15,41^ This is in contrast to other non-haem Fe(II) dioxygenases, including the metazoan oxygen-sensing enzyme hypoxia-inducible factor (HIF) hydroxylases^48^, which typically coordinate Fe(II) with a His/Asp/His-motif.^42,49^ We and others have previously replaced *At*PCO4(−1) His164 with an Asp to mimic this motif, however this completely abolished *At*PCO4 activity.^15,41^ To test whether similar replacement of the other His-triad residues in *At*PCO4 with an Asp (*At*PCO4 H98D and H100D) also resulted in enzyme inactivation, we generated recombinant forms of these variants and tested their capability to catalyse oxidation of RAP2.12_2-15_, an Nt-Cys2 initiating peptidic form of an ERFVII substrate. As anticipated, these mutations also resulted in abolished activity of *At*PCO4 enzymes, likely due to reduced iron retention capability (Figure S3) in line with previous data^15^. Similar results have been reported for replacing the His-triad residues with an Asp in *Hs*CDO-1 in density functional theory studies.^50^ As expected, all residues of the His-triad appear to be essential for the efficient catalysis of Cys oxidation by thiol dioxygenases.

### Residues involved in substrate binding reduce *At*PCO4 activity by decreasing enzyme affinity for peptide substrates

To date, no substrate bound PCO structure has been reported. However, models of PCO^15,41^ and the mammalian thiol dioxygenase ADO^45,51^ in complex with substrates, as well as structures of enzyme-substrate analogue complexes^43^ indicate that substrate binds in the previously predicted region.^42^ Comparison of *At*PCO4 with the structure of *Hs*ADO, which also catalyses oxidation of the Nt-Cys of target proteins^52^, shows that residues Y73 and Y183 are conserved at the potential substrate entry site (Figure S1C, residues in blue); modelled structures of *At*PCO4 docked with peptide substrates suggest that these residues might be important for correct positioning of the Nt-Cys substrate (Figure S5). In previous work investigating the active site residues of *At*PCO4, we reported reduced (but not abolished) activity for a Tyr to Phe variant, *At*PCO4(−1) Y182F^15^. We therefore hypothesised that this and other mutations in this region may decrease PCO activity through reduced substrate binding. This has been confirmed recently for at least the Tyr in *Hs*ADO^43^ corresponding to Y183 in *At*PCO4. We therefore generated and kinetically investigated the *At*PCO4 Y183F variant (equivalent to the *At*PCO4(−1) Y182F variant previously reported^15^) to determine the importance of the Tyr-OH group in substrate binding. We also designed an *At*PCO4 Y73R variant; in *Hs*ADO the equivalent Tyr residue is predicted to form an H-bond with its substrate (Figure S4A) in a docking model and, although this is not predicted for the *At*PCO4-RAP2.12_2-7_ interaction (Figure S4B), we nevertheless hypothesised that a significant change in structure of the residue (introduction of bulky positively charged Arg) at this position may disrupt substrate binding. It should be noted that while this manuscript was in preparation, it was revealed that mutating the equivalent Tyr residue in *Hs*ADO to an Ala or Phe did reduce activity by ∼30 – 70% but did not considerably alter the enzyme’s substrate affinity^43^.

Recombinant *At*PCO4 variants Y73R and Y183F were successfully produced (Figures S6 – S8) and kinetic assays revealed they were both significantly less active towards the peptide substrate RAP2.12_2-15_ than the WT enzyme (Figure 1C) with significantly reduced maximal velocity (*V_max_* of WT ∼2.5, Y73R ∼0.04 and Y183F ∼1.2 μmol.mg^-1^.min^-1^), catalytic constant (*k_cat_* of WT ∼70, Y73R ∼1 and Y183F ∼30 min^-1^) and catalytic efficiency (*k_cat_*/*K_M_* of WT ∼0.69, Y73R and Y183F ∼0.02 min^-1^.μM^-1^, Figure 1D, F, G, Table S5). When a Trp fluorescence-based assay was used to measure binding affinity of the enzymes for substrate,^53^ the results revealed that the decrease in activity could be ascribed to a considerably weakened interaction between the variants and RAP2.12_2-15_ (Figure 1H, *K_d_* of WT ∼200, Y73R ∼3000 and Y183F ∼3500 μM). Iron retention capacity was not reduced in either variant (iron occupancy of WT ∼17%, Y73R ∼33% and Y183F ∼50%; Figure S9B). Overall, the data indicated that *At*PCO4 residues Y73 and Y183 play an important role in substrate binding and as a consequence impact effective catalysis. While removal of the Y183-OH group resulted in reduced *At*PCO4 activity, the substitution Y73R resulted in an almost complete loss of enzymatic activity.

**Figure 1.**
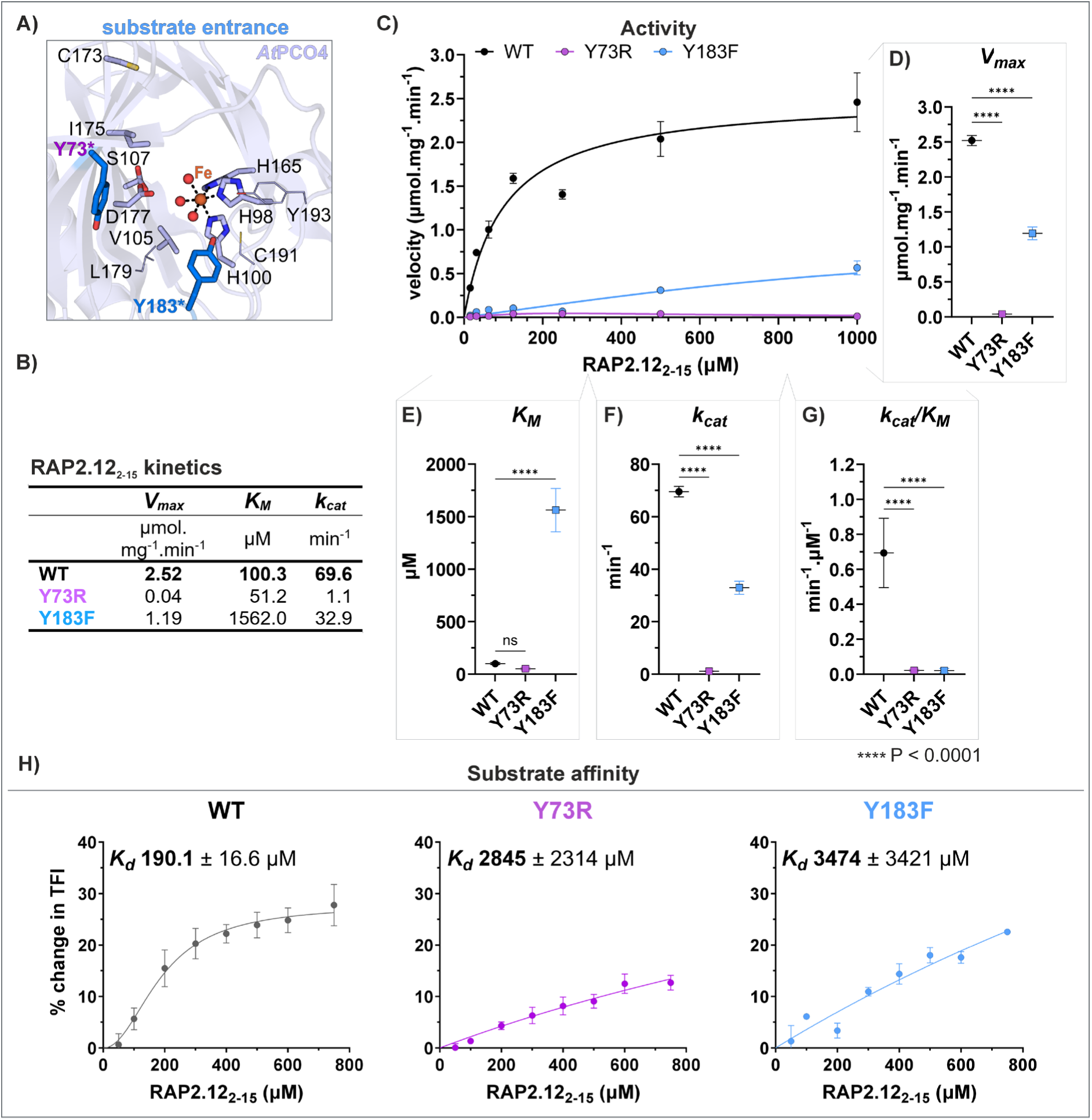
Activity and substrate binding of substrate entry site variants. **(A)** Cartoon presentation of *At*PCO4 crystal structure (PDB: 6S7E, crystal structure of *At*PCO4-1 but residues were numbered according to *At*PCO4-2 to align with main text) with investigated residues highlighted in bright blue and with asterix. **(B)** Values of the Michaelis-Menten kinetics for *At*PCO4 enzymes with substrate peptide. Parameters for Y183F were determined based on additional data shown in the supplementary (Figure S9C). **(C)** Initial velocities of *At*PCO4 WT (from two independent assays in technical triplicates, 2 x n = 3) and variants (in technical triplicates, n = 3) from 0 – 2 min time courses plotted over peptide substrate concentrations. Standard deviation is indicated by error bars. **(D)** – **(G)** Graphic overview of **(D)** maximal velocity, **(E)** Michaelis-Menten constant, **(F)** catalytic constant and **(G)** catalytic efficiency of *At*PCO4 WT and variant values in **(B)** calculated using the Michaelis-Menten equation in GraphPad Prism, Version 10.1.2. Error bars indicate the standard error of the mean. The statistical significance (P < 0.05) of the kinetic constants from each variant in comparison to *At*PCO4 WT were evaluated using one-way ANOVA followed by Holm-Šídák’s test to correct for multiple comparisons using hypothesis testing (GraphPad Prism). **(H)** Affinity of RAP2.12_2–15_ to *At*PCO4 enzymes determined using an intrinsic fluorescence tryptophan assay.

### Residues close to active site Fe(II) alter catalysis independent of overall substrate affinity

We next wanted to explore the effects of mutating residues that might have a role in *At*PCO4 catalysis. These are likely to be residues close to the active site Fe(II) and potentially in a position to interact with the Nt-Cys, O_2_ and/or reaction intermediates. The carboxylate group of residue D177 is expected to stabilise the Nt-Cys amine or thiol of protein substrates (D176 in *At*PCO4-1)^15,41^ similar to the mammalian ADO (D206 in *Hs*ADO)^51^; this was also seen in *in silico* docking models (Figure S4). Removing the negative charge of the Asp residue by replacing it with an Asn (*At*PCO4(−1) D176N) has previously been shown to significantly reduce the activity of *At*PCO4.^15^ We therefore reasoned that substituting the Asp with a Glu (D177E) should still reduce *At*PCO4 activity but to a lesser degree than substituting it with an Asn (as previously reported^15^) because the negative charge would be maintained. During the preparation of this manuscript the equivalent Asp to Glu mutation was examined in the human homologue ADO resulting in ∼20% activity compared to WT while Asp to Asn mutation resulted in ∼2% activity^43^. *At*PCO4 residues V105 and S107, located approximately *trans* to the His-triad of *At*PCO4 (Figure 2A, residues in green), were replaced with Gly and Leu, respectively, representing their equivalent residues in *Hs*ADO (Figure S1C). Since *Hs*ADO has a high *K_M_*(O_2_) (∼55 μM^52^), and due to the position of residues V105 and S107, it was hypothesised that these residues might affect the positioning of O_2_ and Nt-Cys substrate or reaction intermediates. While an *At*PCO4 S107A variant previously showed a similar activity to *At*PCO4 WT,^15^ a Leu substitution would be more bulky and thereby have a stronger impact on *At*PCO4 activity.

**Figure 2.**
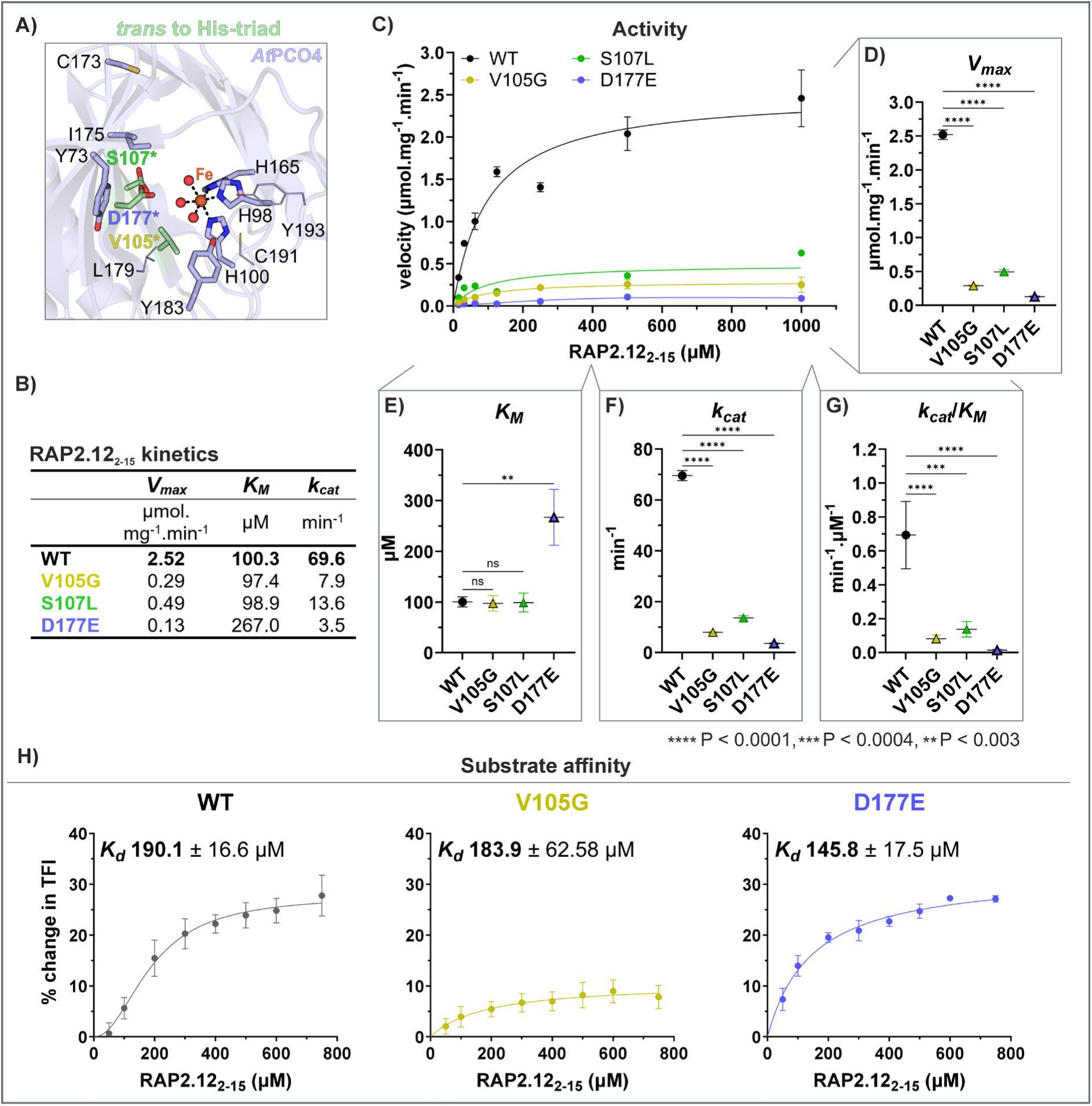
Activity and substrate binding of variants with mutations approximately *trans* to the His-triad. **(A)** Cartoon presentation of *At*PCO4 crystal structure (PDB: 6S7E, crystal structure of *At*PCO4-1 but residues were numbered according to *At*PCO4-2 to align with main text) with investigated residues highlighted in green and with asterix. **(B)** Values of the Michaelis-Menten kinetics for *At*PCO4 enzymes with substrate peptide. **(C)** Initial velocities of *At*PCO4 WT (from two independent assays in technical triplicates, 2 x n = 3) and variants (in technical triplicates, n = 3) from 0 – 2 min time courses plotted over peptide substrate concentrations. Standard deviation is indicated by error bars. **(D) – (G)** Graphic overview of **(D)** maximal velocity, **(E)** Michaelis-Menten constant, **(F)** catalytic constant and **(G)** catalytic efficiency of *At*PCO4 WT and variants calculated using the Michaelis-Menten equation in GraphPad Prism. Error bars indicate the standard error of the mean. The statistical significance (P < 0.05) of the kinetic constants from each variant in comparison to *At*PCO4 WT were evaluated using one-way ANOVA followed by Holm-Šídák’s test to correct for multiple comparisons using hypothesis testing (GraphPad Prism). Only significant differences to *At*PCO4 WT are displayed. **(H)** Affinity of RAP2.12_2–15_ to *At*PCO4 enzymes determined using an intrinsic fluorescence tryptophan assay^53^. The *K_d_* was estimated using the one-site or Hill-slope specific binding model in GraphPad Prism. The % change in tryptophan fluorescence intensity (TFI) was lower in *At*PCO4 V105G compared to the WT enzyme which might reflect a change in the microenvironment of the Trp residue due to mutation V105G that did not affect substrate binding.

Recombinant *At*PCO4 variants D177E, V105G and S107L were produced (Figure S6 – 8) and kinetic analysis revealed that all mutations decreased *At*PCO4 activity compared to WT enzyme, demonstrated by significantly lower *V_max_* (WT ∼2.5, V105G ∼0.3, S107L ∼0.5 and D177E ∼0.1 μmol.mg^-1^.min^-1^), *k_cat_* (WT ∼70, V105G ∼8, S107L ∼14 and D177E ∼4 min^-1^) and catalytic efficiency values (WT ∼0.69, V105G ∼0.08, S107L ∼0.14 and D177E ∼0.01 min^-1^.μM^-1^; Figure 2B – G, Table S5). Interestingly, variants V105G and S107L showed similar *K_M_* values to WT (*K_M_*of WT ∼100, V105G ∼97 and S107L ∼99 μM; Figure 2B and E), indicative that these mutations did not affect substrate binding. This was confirmed in a binding assay for variant V105G with a *K_d_*(V105G) of ∼184 ± 63 μM that was similar to the *K_d_*(WT) ∼190 ± 17 μM of the WT enzyme (Figure 2H). The lower iron occupancy of variant S107L may contribute to the lower activity of this variant (Figure S10B). Our binding assay revealed that variant D177E also had a slightly reduced *K_d_* value compared to the WT enzyme (*K_d_* of WT ∼190 ± 17 μM and D177E ∼150 ± 18 μM; Figure 2H) suggesting similar or higher affinity of variant D177E to RAP2.12_2-15_ substrate. The *K_M_* value of variant D177E (∼270 μM) was likely high (relative to the WT) due to the low activity (*k_cat_*) of this variant (Figure 2B and F). *In silico* docking models indicated that a Glu residue at position 177 can either coordinate the iron or still coordinate the substrate (Figure S10D), both of which could affect the positioning of the iron and the substrate in the active site, consistent with the D177E mutation altering the rate of catalysis rather than reducing substrate or iron binding.

We next investigated residues C173 and I175, which are positioned approximately *trans* to the substrate entry site and hypothesised that they might have a role in O_2_ interactions and/or facilitating steps in catalysis (Figure 3A, residues in magenta). Interestingly, residue C173 is conserved in PCOs but not in *Hs*ADO nor *Hs*CDO-1 which both comprise an Ala residue at this position (A202 in *Hs*ADO and A151 in *Hs*CDO-1, Figure S1C). Residue C173 was therefore mutated to an Ala. The equivalent residue to *At*PCO4 I175 in *Hs*ADO is a Phe, while in *Hs*CDO-1 it is a Ser (Figure S1C, residues in magenta). The bulkier Phe in *Hs*ADO might restrict movement of O_2_ around the iron and, therefore, also O_2_ activation, consistent with the higher sensitivity of *Hs*ADO to changes in O_2_ concentrations than *At*PCO4. *At*PCO4 I175 was therefore mutated to a Phe. Recombinant *At*PCO4 variants C173A and I175F were successfully produced (Figures S6 – S7) and kinetically analysed.

**Figure 3.**
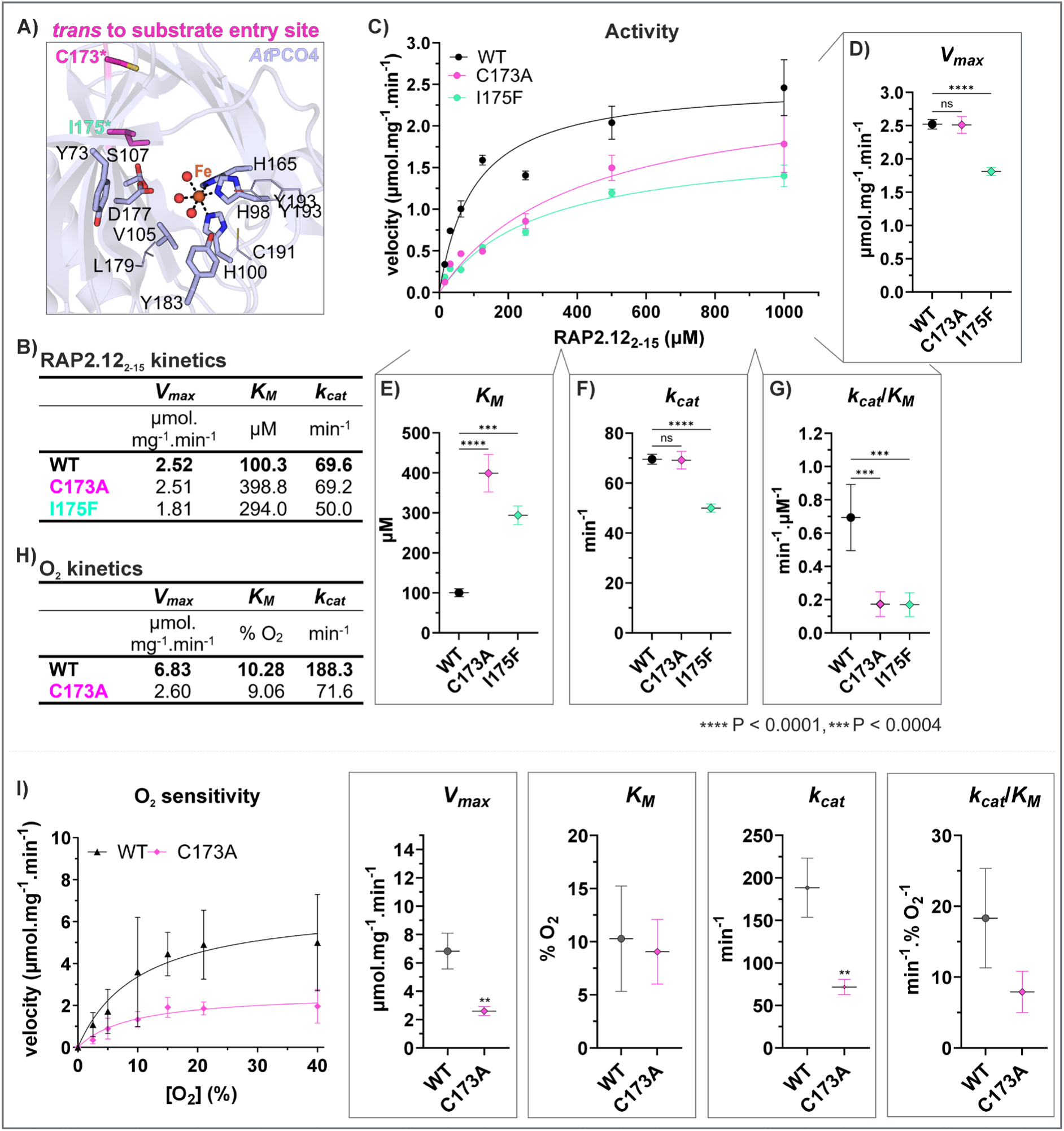
Activity and substrate binding of residues affecting *At*PCO4 catalysis and being located approximately *trans* to the substrate entry site. **(A)** Cartoon presentation of *At*PCO4 crystal structure (PDB: 6S7E, crystal structure of *At*PCO4-1 but residues were numbered according to *At*PCO4-2 to align with main text) with investigated residues highlighted in magenta and with asterix. **(B)** Values of the Michaelis-Menten kinetics for *At*PCO4 enzymes with substrate peptide. **(C)** Initial velocities of *At*PCO4 WT (from two independent assays in technical triplicates, 2 x n = 3) and variants (in technical triplicates, n = 3) from 0 – 2 min time courses plotted over peptide substrate concentrations. Standard deviation is indicated by error bars. **(D)** – **(G)** Graphic overview of **(D)** maximal velocity, **(E)** Michaelis-Menten constant, **(F)** catalytic constant and **(G)** catalytic efficiency of *At*PCO4 WT and variants calculated using the Michaelis-Menten equation in GraphPad Prism. Error bars indicate the standard error of the mean. The statistical significance (P < 0.05) of the kinetic constants from each variant in comparison to *At*PCO4 WT were evaluated using the Michaelis-Menten model. Standard deviation is indicated by error bars. The statistical significance (P < 0.05) of kinetic parameters was evaluated as for **(D)** – **(G)** (**: P = 0.0013, GraphPad Prism).

*At*PCO4 I175F demonstrated reduced catalytic efficiency (*k_cat_*/*K_M_*of I175F ∼0.2 min^-1^.μM^-1^ compared to WT ∼0.7 min^-1^.μM^-1^) by decreasing the catalytic turnover rate (*k_cat_* of I175F ∼50 min^-1^ compared to WT ∼70 min^-1^) and increasing the *K_M_* of *At*PCO4 (*K_M_* of I175F ∼290 μM compared to WT ∼100 μM; Figure 3B, E – G, Table S5). Variant C173A demonstrated *V_max_* and *k_cat_* values similar to the WT enzyme (*V_max_*∼2.5 μmol.mg^-1^.min^-1^ and *k_cat_* ∼70 min^-1^), while its catalytic efficiency was decreased (*k_cat_*/*K_M_*(WT) ∼0.7 min^-1^.μM^-1^, *k_cat_*/*K_M_*(C173A) ∼0.2 min^-1^.μM^-1^) owing to a higher *K_M_* value (*K_M_*(WT) ∼100 μM, *K_M_*(C173A) ∼400 μM; Figure 3B – G).

The location of residues C173 and I175 approximately *trans* to the substrate entry site and > 6.5 Å distant from the Nt-Cys substrate (Figure S5) indicated that the higher *K_M_*values observed for these variants may reflect different requirements for the O_2_ co-substrate rather than the peptide substrate RAP2.12_2-15_. Focussing on the C173A variant, due to its apparent similar *V_max_* to *At*PCO4 WT (Figure 3B – D), we therefore determined the kinetic parameters of this variant with respect to O_2_, under conditions of optimal RAP2.12_2-15_ availability. Measuring the activity of *At*PCO4 C173A at 0 to 40% O_2_ with 1500 μM RAP2.12_2-15_ revealed that variant C173A could not use O_2_ as efficiently as the WT enzyme (Figure 3I). This was represented by a *K_M_*(O_2_) similar to the WT enzyme (*K_M_*(O_2_) ∼10 % O_2_) but lower *V_max_* (*V_max_*(O_2_) of WT ∼7 μmol.mg^-1^.min^-1^ and C173A ∼3 μmol.mg^-1^.min^-1^) and *k_cat_* values (*k_cat_*(O_2_) of WT ∼190 min^-1^ and C173A ∼70 min^-1^; Figure 3H and I). We noted that in the O_2_ assay, WT enzyme was able to achieve a greater *V_max_* than in the RAP2.12_2-15_ assay, potentially due to limiting O_2_ in normoxic experiments as well as differences in assay conditions (Figure S11C). To confirm that mutating a residue that is located approximately *trans* to the substrate entry site mainly affects the catalysis of the dioxygenation of substrate peptide and not peptide substrate binding, the *K_d_*of variant C173A was determined (Figure S11E). The *K_d_* of variant C173A was similar to the WT enzyme (*K_d_*(WT) ∼190 ± 17 μM, *K_d_*(C173A) ∼250 ± 52 μM, Figure S11D and E) and also *in silico* docking models showed Nt-Cys positioning in variant C173A similar to that in the WT enzyme (Figure S11A). Overall, variant C173A therefore was found to catalyse the dioxygenation of RAP2.12_2-15_ peptide slower than the WT enzyme but was the most active of all the variants studied.

### Reduced *At*PCO4 activity has the potential to increase submergence tolerance in *4pco*

#### Arabidopsis plants while not significantly altering the phenotype in normoxia

We have previously reported that *At*PCO4 variants with biochemically ablated activity, when introduced into the *4pco* Arabidopsis model, also showed ablated activity *in vivo.*^15^ To improve submergence tolerance in plants, our strategy seeks to enhance ERFVII stability and thereby function during a submergence event, through reduced *At*PCO activity, while maintaining *At*PCO-initiated degradation of ERFVIIs under normoxic conditions. We therefore selected two *At*PCO4 variants that had shown reduced or similar activity to *At*PCO4 WT *in vitro*, to introduce into Arabidopsis *4pco* plants and test their impact *in vivo*. We selected *At*PCO4 C173A as it demonstrated good catalytic capability in normoxia, but limited activity compared to WT enzyme in response to O_2_ availability. We also selected variant Y183F to test the effect of significantly reduced substrate binding on *At*PCO4 function *in vivo*. Although we were not able to determine kinetic parameters of the Y183F variant with respect to O_2_, as it was not possible to reach saturating RAP2.12_2-15_ substrate conditions, we did find that this variant showed sensitivity to O_2_ availability (Figure S9C).

As previously reported, *4pco* plants, which lack all *At*PCOs except *At*PCO3, exhibit developmental defects that can be complemented with functional *At*PCO enzymes to establish a Col-0 like phenotype.^15,16,52^ We therefore established two lines per genotype for *4pco AtPCO4* WT (wt-1 and wt-2) and for *4pco AtPCO4-C173A* (C173A-1 and C173A-2); we were only able to establish one line for *4pco AtPCO4-Y183F* (Figure S12A, sequences in Figures S13 – S15).

The complemented *4pco* lines had a phenotype similar to that of Col-0 plants (Figure S12B). This is in contrast to the phenotype we previously reported for *4pco* plants complemented with inactive *At*PCO variants^15^ which showed a phenotype more similar to that of *4pco* plants^52^. Nevertheless, small differences were noticed between the lines: wt-1 and C173A-2 plants germinated and grew at a rate similar to Col-0, whereas wt-2 and Y183F were slightly slower followed by C173A-1 (Figure 4A: ‘before’ panel, Figure S12B).

**Figure 4.**
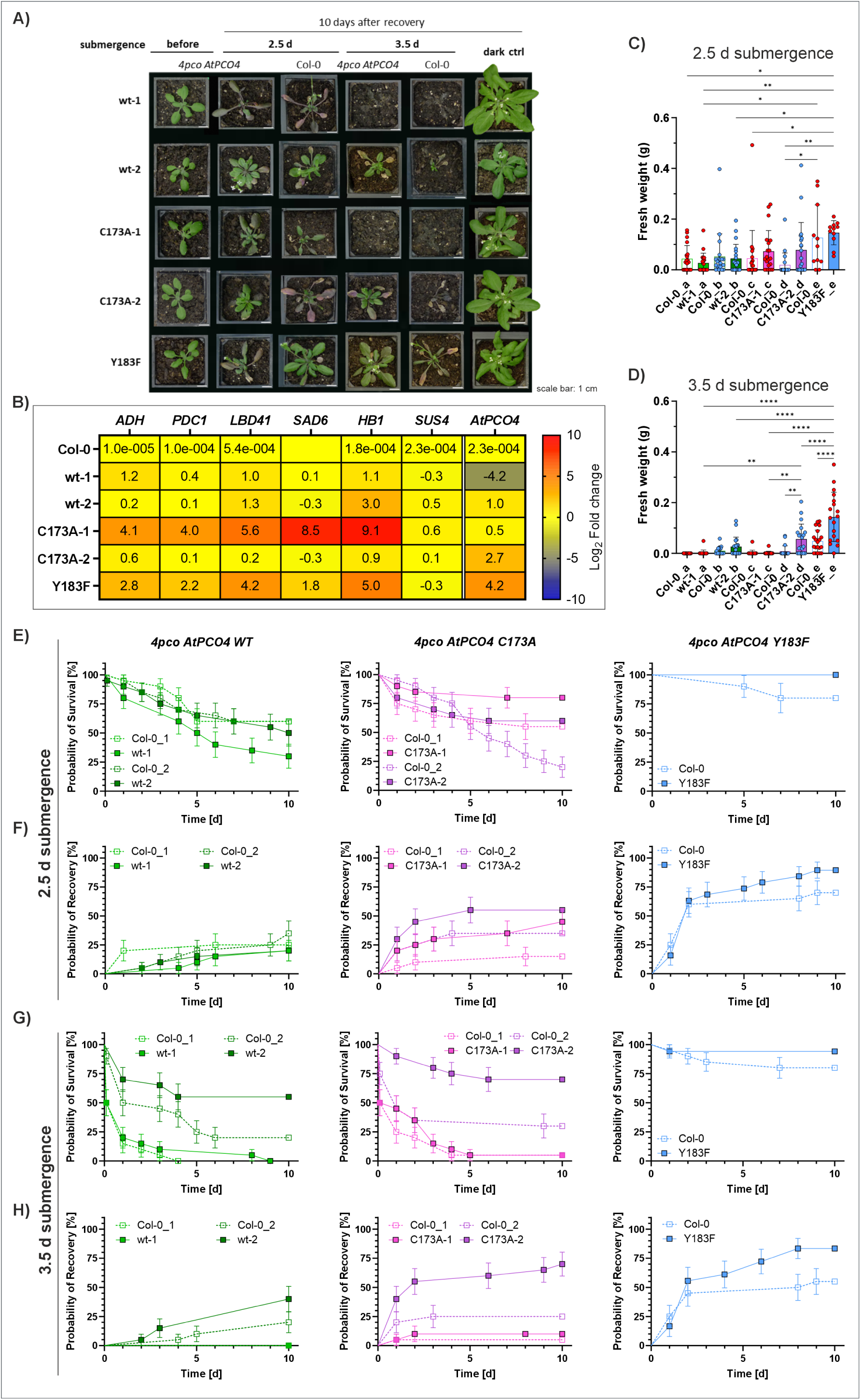
Effect of *At*PCO4 variants in the *4pco AtPCO4* Arabidopsis plant model. **(A)** Panel of submerged plants from the second experiment showing the most recovered plant out of the 10 submerged plants from each genotype. **(B)** Change in transcript levels of HRGs and *AtPCO4* WT and mutants (primer pair: qpAtPCO4-2) in 5.5-week-old *4pco AtPCO4* and Col-0 plants. Two independent experiments with two technical replicates of four biological samples each (n = 2 x (2 x 4)). **(C)** and **(D)** fresh weight of plants submerged for **(C)** 2.5 and **(D)** 3.5 days after 10 days of recovery (lower case letters indicate plants submerged in the same box). Dead plants were considered to weigh 0 g. The statistical significance (P < 0.05) was determined using two-way ANOVA followed by Tukey’s multiple comparisons test (GraphPad Prism). Only significant differences between *4pco AtPCO4* lines and their control Col-0 lines or other *4pco AtPCO4* lines are displayed (**: P < 0.0084; ****: P < 0.0001). **(E)** and **(G)** survival of each *4pco AtPCO4* line (continuous line) and its Col-0 control (dashed line, submerged in same box) after **(E)** 2.5 and **(G)** 3.5 days submergence was scored using a dead/alive scoring system. To be considered alive, the plant needed to have a green meristem with at least the area of one leaf in a fresh green colour attached. Values represent two independent experiments with 10 plants / line / experiment (n = 2 x 10), except for line Y183F (2.5 days submergence: 10 plants, 3.5 days submergence: n = 8 + 10 plants). **(F)** and **(H)** recovery rate of each *4pco AtPCO4* line and its Col-0 control group after **(F)** 2.5 and **(H)** 3.5 days submergence is shown as described for **(E)** and **(G)**. A plant was scored as recovering, when it produced a new leaf and that leaf continued to grow. Abbreviations: ADH: alcohol dehydrogenase, PDC1: pyruvate decarboxylase 1, LBD41: LOB domain-containing protein 41, SAD6: stearoyl-acyl Carrier Protein 9-Desaturase 6, HB1: plant haemoglobin 1, SUS4: sucrose synthase 4, *AtPCO4*: *AtPCO4* insert.

Before examining the impact of the mutations on HRG transcription and submergence recovery, we checked the transcript level of each introduced *AtPCO4* gene as both its transcript level and its catalytic capability can affect ERFVII protein levels and consequent effects on HRGs. We compared *AtPCO4* transcription to the housekeeping gene *Polyubiquitin 10* (*UBQ10*), normalised to Col-0, and found that transcription levels varied amongst the different lines (Figure 4B): wt-1 < Col-0 < C173A-1 < wt-2 < C173A-2 < Y183F (Figure 4B, Figure S16). The apparent downregulation of *AtPCO4* transcription in wt-1 was surprising since its phenotype was similar to that of Col-0 (Figure S12B). It is possible that the high catalytic efficiency of *At*PCO4^46^ results in similar levels of ERFVII oxidation to Col-0 despite lower expression levels.

We next measured the levels of HRG transcripts in each of the *4pco AtPCO4* mutant lines relative to housekeeping gene *UBQ10* (Figure 4B). All of the tested marker genes for hypoxia responses (except *SUS4* encoding sucrose synthase 4) showed increased transcript levels for line C173A-1 and to a lesser degree for line Y183F, while their transcript levels in line C173A-2 were similar to wt-1, wt-2 and Col-0 (Figure 4B, Figure S17). Collectively, the increased HRG levels in the variant lines (at *AtPCO4* transcript levels similar to WT) suggested that the *AtPCO4* mutations reduced the activity of *At*PCO4 *in vivo* leading to ERFVII stabilisation. However, the effect of decreased *At*PCO4 activity could be, in part, neutralised by higher *AtPCO4* mutant transcription, as seen in the case of line C173A-1 vs line C173A-2 (Figure 4B). The increased *ADH*, *PDC1* and *LBD41* transcript levels observed for line Y183F, despite high *AtPCO4 Y183F* transcription, likely arise due to the significantly lower activity of *At*PCO4 Y183F compared to C173A *in vitro*. Overall, the elevated HRG transcript levels for lines C173A-1 and Y183F in normoxia indicated that the variants were less active than *At*PCO4 WT *in vivo*. However, the phenotype of the *4pco At*PCO4 variant plants was more similar to *4pco AtPCO4* WT plants (Figure 4A, ‘before’ and ‘dark ctrl’ panels) than *4pco* plants^52^, implying that the introduced *At*PCO4 variants were at least partially active *in vivo* in normoxia. Overall, the relative kinetic properties of the *At*PCO4 variants appeared to be consistent between *in vitro* and *in vivo* conditions.

To assess if the C173A and Y183F variants could improve submergence tolerance compared to *At*PCO4 WT, ten plants of each line were submerged in the dark for 2.5 or 3.5 days alongside Col-0 plants as controls and dark-only (not submerged) controls. To minimise the effect of different survival and recovery due to plant age^54^, plants with a 3-week Col-0 phenotype were chosen for the submergence experiment. The survival and recovery rates after submergence were evaluated during a 10-day recovery period (Figure 4E – H). The results of the 2.5 days submergence experiment showed that more plants containing either *At*PCO4 C173A or Y183F survived and recovered than their Col-0 control plants (Figure 4E and F, Table S6, Figure S18, left panel). In contrast, fewer plants of wt-1 and wt-2 survived and recovered compared to their Col-0 control plants (Figure 4E and F, Table S6). The fresh weight of 2.5 days submerged plants after 10-days of recovery was similar for the variant lines, C173A-1, C173A-2 and Y183F, with plants from line Y183F being significantly heavier than wt-1 and wt-2 (Figure 4C, for dry weight and weight of dark control plants see Figure S19A – D). However, the Col-0 control plants for line Y183F were also heavier than other Col-0 plants (Figure 4C). The differences in fresh weight were more substantial for plants recovering from 3.5 days submergence (Figure 4D, for dry weight and weight of dark control plants see Figure S19E – H). The fresh weight of recovered plants from lines, C173A-2 and Y183F, was significantly higher than of the other lines, especially for plants from line Y183F (Figure 4D). Nevertheless, the occurrence of zero survival of wt-1 + Col-0 and C173A-1 + Col-0 in at least one of the two experimental repeats (Figure S18, right panel), affected this statistic as well as the reduced survival rate for both lines, wt-1 and C173A-1 (Figure 4G). While the survival and recovery rates of wt-1 and C173A-1 were similar to their respective Col-0 plants, lines C173A-2 and Y183F appeared to survive and recover better after 3.5 days submergence than their Col-0 controls (Figure 4G and H, Figure S18, right panel). Interestingly, a greater number of plants from line wt-2 survived (compared to its Col-0 control) after 3.5 days submergence (Figure 4G), however, the surviving plants did not seem to grow and gain as much biomass as plants with Y183F (Figure 4D, Figure S18, right panel). A possible explanation could be that the lack of *At*PCO1, 2 and 5 in complemented *4pco* plants resulted in a prolonged stabilisation of ERFVIIs despite the re-introduced *At*PCO4 WT enzyme compared to Col-0. However, this stabilisation seemed to be insufficient to promote biomass gain to the same extent as *At*PCO4-Y183F (Figure 4D).

In summary, the potential for increasing HRG transcript levels in *4pco AtPCO4* variants lines, C173A-1, −2 and Y183F, compared to wt-1 and wt-2 correlated with the reduced activity of the *At*PCO4 variants observed *in vitro*. The capability to increase stabilisation of ERFVIIs and thereby transcription of HRGs appeared to be beneficial for submergence tolerance, however the results also demonstrated the importance of ERFVII destabilisation during normoxia (line C173A-1). Overall, *At*PCO4 variants with reduced activity could improve the recovery and survival of complemented *4pco* Arabidopsis plants subjected to submergence, especially when the HRGs were not induced in normoxic conditions. This suggests that fine-tuning PCO activity could indeed be used to improve flood tolerance.

## Discussion

PCOs are critical oxygen-sensing enzymes that catalyse the first step in destabilising ERFVII transcription factors.^9,12^ Their reduced activity at low O_2_ concentrations stabilises ERFVIIs to upregulate genes that enable acclimation to hypoxic conditions.^7–9^ This is an important element for plants to be able to survive temporary periods of submergence. As climate change causes an increase in the intensity and duration of flood events, it is important for food security that crop plants are sufficiently resilient to this stress.^55^ Elevated stabilisation of ERFVIIs has been associated with improved submergence resilience.^8,37^ We have proposed that reducing the activity of PCOs could be a mechanism to achieve this, particularly if their activity can be controlled such that ERFVII levels remain low in normoxia but stabilise earlier and destabilise later over the course of a hypoxic event such as a flood.^2,40^

In this study, we reduced *At*PCO4 activity first *in vitro* and then *in vivo* by mutating residues localised around the active site using a structure-guided approach. The mutations affected iron binding, substrate binding and enzymatic catalysis. Kinetic and biophysical analysis of variants at selected residues confirmed their predicted contribution to ERFVII Nt-Cys oxidation. *At*PCO4 variants of iron-coordinating residues, H98D and H100D, showed ablated activity, likely due to distorted iron binding capacity. *At*PCO4 Y73R also showed ablated activity, though this was due to severely weakened substrate binding. The more structurally conservative *At*PCO4 Y183F variant at the substrate binding site also caused weakened substrate affinity and consequently reduced (but not ablated) activity. Mutations of residues closer to the catalytic site of *At*PCO4, D177E, V105G and S107L, reduced catalytic rate and efficiency but had no substantial impact on substrate affinity. Variant D177E is likely still able to coordinate the substrate Nt-Cys thiol or amine group via its carboxy group as suggested for its human homologue ADO^43^. The severely reduced activity of this variant might be due to the larger residue’s capability to mis-position the iron centre, Nt-Cys, O_2_ or reaction intermediates. Variant S107L likely reduced iron binding capacity and could have disturbed the positioning of O_2_, reaction intermediates and/or the *At*PCO4 D177-carboxylate group. Residue V105 might be involved in optimising the position of the O_2_ co-substrate in the active site by limiting the space elsewhere. Variants of residues located *trans* to the substrate entry site, C173A and I175F, might have reduced the catalytic efficiency of *At*PCO4 by altering the position, and therefore the stability of O_2_/reaction intermediates. Mutation C173A decreased the enzyme’s catalytic rate once O_2_ was coordinated in the active site, though whether this effect is through direct interactions or conformational changes remains to be elucidated. In a recent crystal structure of the human homologue ADO with a bound substrate analogue (PDB: 9DXB) both residues, C173 and I175 (*At*PCO4 numbering), would be too distant from the reaction centre to have a direct impact on catalysis. Our biochemical data were consistent with similar studies from other thiol dioxygenases, a detailed discussion of which can be found in the Supplementary Information file.

We selected two mutations that reduced the activity of *At*PCO4 *in vitro* to complement a *4pco* Arabidopsis plant model to investigate whether a decrease in *At*PCO4 activity could improve submergence tolerance. We selected one variant with a severe effect on *in vitro At*PCO4 activity, Y183F, which caused a reduction in substrate affinity, and one variant with a relatively mild effect on *At*PCO4 activity, C173A, which caused a reduction in catalytic turnover. Both variants resulted in improved recovery after submergence following 2.5 and particularly 3.5 days of dark submergence. Reduced *At*PCO4 activity correlated with a ∼130 – 300% increase in recovery and ∼100 – 300% increase in survival probability compared to ∼60 – 200% and ∼50 – 280% for *At*PCO4-2 WT, respectively (both in relation to Col-0). While it is hard to dissect at which point during the submergence regime the decreased *At*PCO4 activity conferred the greatest advantage, the improved recovery implied that reduced *At*PCO4 activity was particularly beneficial during/after de-submergence. In this case, the reduced activity of *At*PCO4 C173A or Y183F could have provided plants with more time to adapt to the sudden increase in O_2_ upon de-submergence. *At*PCO4 transcript levels also appeared to be important in controlling HRG transcript levels and submergence tolerance, indicating that both the absolute amount of enzyme as well as its intrinsic activity are important in determining ERFVII stability and HRG transcription. This is in accordance with previous data showing that *At*PCO4 is important for submergence tolerance.^16^ Notably, reduced *At*PCO4 activity increased submergence tolerance to a greater degree when HRGs were not upregulated during normoxia, as was the case for C173A-2 compared to C173A-1, in accordance with previously published results that temporary but not constitutively stabilised ERFVIIs are able to increase submergence tolerance.^8,56^ When ERFVIIs are stabilised during normoxia they promote fermentative metabolism, e.g., by inducing the transcription of *ADH1* and *PDC1*, which consumes carbohydrate resources, thus, fewer resources will be available for the plant during submergence leading to lower submergence tolerance.^8,16,56^ This additional stress could potentially be observed in plants from line C173A-1 and resulted in reduced survival and recovery compared to line C173A-2. However, the submergence tolerance of C173A-1 was still improved after 2.5 days submergence compared to lines with *At*PCO4 WT.

Overall, our results demonstrate that engineered *At*PCO4 with lower activity has indeed the potential to upregulate HRG transcript levels and increase the submergence tolerance of plants. Although our experiments indicate that both absolute transcription levels and the catalytic ability of mutant PCOs are important in conferring improved submergence tolerance, ultimately editing genomic PCO sequences to introduce key mutations could negate the requirement to control for PCO transcript levels. Our study is a starting point to further engineer PCOs with more finely-tuned activities to balance stabilisation of ERFVIIs at the onset/offset of hypoxia and destabilisation during normoxia to avoid impacting yield. The high conservation of active site residues within the PCOs of flowering plants^16^ means that the strategy can likely be applied to other PCOs in Arabidopsis but also agronomically relevant species; the occurrence of multiple PCO isoforms in many agronomically relevant species suggests the potential to fine-tune cultivars’ submergence tolerance and growth. In crops that are amenable to CRISPR-Cas gene editing methods, it may be feasible to introduce single amino acid substitutions to PCOs; a similar approach has been used to increase the resistance of tomatoes to late blight by altering an endogenous immune protease.^57,58^ Additionally, tailoring the activity of PCOs could provide an advantage to other abiotic stresses like heat and salt stress as the Cys/Arg-N-degron pathway has been connected to both.^59,60^ Ultimately, using an enzyme engineering approach to design, test and introduce targeted mutations into PCOs of agronomically relevant species could contribute to future food security amidst extreme weather caused by climate change.

### Experimental Procedures

#### Site-directed mutagenesis and protein production

The thrombin cleavage site in pET28a(+)-His-thrombin-AtPCO4-1 from White *et al.* 2018 was exchanged to a TEV cleavage site by Genscript using the *NcoI* and *XhoI* restriction sites. Specific sites in pET-28a(+)-His-TEV-AtPCO4-1 (recombinant protein synthesis, deposited at addgene #191802) and pENTR_PCO4_AtPCO4 were mutated using the QuikChange® Site-Directed Mutagenesis Kit (Stratagene) protocol. Primers were designed according to Zheng, Baumann and Reymond^61^ and are listed in Supplementary Table S1. After confirming successful mutants by sequencing, proteins from pET28a(+) plasmids were overexpressed in chemically competent *Escherichia coli* (*E. coli*) BL21 (DE3) cells and purified as described previously^46,47^ with some adaptations. In short, the filtered supernatant was purified using Ni(II)-affinity chromatography. The *At*PCO4 proteins were eluted during an imidazole concentration from 30 – 100%. After removal of imidazole using a desalting PD10 column, the His-tag was cleaved by incubating the proteins with TEV protease (purified from plasmid pRK793^62^ (addgene)) for 14 – 16 hours. Cleaved protein was separated from uncleaved protein, His-tags and TEV protease via reverse Ni(II)-affinity chromatography. The untagged *At*PCO4 proteins were purified further by size exclusion chromatography and aliquots were stored at −80 °C. Protein concentration was determined by UV/Vis-spectroscopy and protein purity was estimated by sodium dodecyl-sulfate polyacrylamide gel electrophoresis (SDS-PAGE).

#### Determination of iron occupancy in *At*PCO4 enzymes

The iron binding capacity of *At*PCO4 variants was established using a colorimetric method with 1,10-phenanthroline and/or inductively coupled plasma mass spectrometry (ICP-MS) as described previously^46,63^. In short, to estimate the iron content using 1,10-phenanthroline, the protein samples were denaturated and the isolated iron was exposed to 1,10-phenathroline. The absorption of the Fe(II)-1,10-phenanthroline complex was measured at 535 nm and the Fe(II) content of each sample was calculated from a standard curve (0 – 250 μM FeSO_4_).

To determine the iron content using ICP-MS, the protein samples were diluted in 2% nitric acid and submitted to an ICP-MS autosampler. The instrument was calibrated with dilutions from two standards. The quality of the detection methods was tested with an external standard. The iron concentration of each sample was measured alongside an internal Rhodium standard to account for any instrumental drift.

#### *At*PCO4 activity assay

Initial activity screens, optimisation of exogenous iron concentrations, kinetic assays and O_2_ sensitivity assays were conducted, and the dioxygenation of the substrates was determined by mass spectrometry as described previously^47^. In short, enzyme master mixes (MM) were prepared in 50 mM bis-tris propane (1,3-bis[tris(hydroxymethyl)methylamino]propane) and 50 mM NaCl in LC/MS grade water and supplemented with FeSO_4_ and (+)-sodium L-ascorbate. Substrate MMs were prepared by resuspending RAP2.12_2-15_ (14-mer) peptide substrate in the same buffer as described above which was supplemented with tris(2-carboxyethyl)phosphine hydrochloride (TCEP) to prevent dimerization. Enzyme and substrate were pre-incubated for 5 – 10 min at 25 °C, before starting the reaction by adding the enzyme MM into the substrate MM. The reaction solution was quenched in 5% (v/v) formic acid at the respective time points and once an assay was finished, it was stored at −20 °C. The samples were analysed using an ultra-performance liquid chromatography (UPLC) system coupled to a quantitative time of flight (Q-TOF) mass spectrometer operated in positive ion/electrospray mode. The source conditions were optimised for minimal fragmentation and maximal sensitivity. Product concentrations were calculated by integrating the area under the chromatogram peaks of the substrate and product (+32 Da) followed by determining the proportion of the product from the total peptide concentration in the sample.

For the O_2_ sensitivity assay, the same principle was applied although using a different setup as previously described in detail^47^.

#### Binding assay using intrinsic tryptophan fluorescence

The affinity of RAP2.12_2-15_ peptide substrates to *At*PCO4 WT and variants was determined using the intrinsic fluorescence of a Trp residue close to the active site (W121 in *At*PCO4 isoform 2) as described previously^53^. In short, iron was removed from *At*PCO4 enzymes to prevent substrate turnover by adding a dialysis step with 10 mM 1,10-phenanthroline and 50 mM ethylenediaminetetraacetic acid (EDTA) after the SEC step during protein purification. Samples were prepared under anaerobic conditions at room temperature. The iron removed enzymes were diluted in anaerobic assay buffer (50 mM Tris, 400 mM NaCl, pH 8) and supplemented with NiCl_2_. RAP2.12_2-15_ peptide substrate was dissolved in assay buffer, supplemented with TCEP, and diluted to the appropriate substrate concentrations. Enzyme and substrate solutions were dispensed into a 96-well black flat-bottom microplate in anaerobic conditions. The intrinsic Trp fluorescence of the enzyme was measured at λ_ex_ / λ_em_ 280/350 nm, atmospheric conditions and 25 °C with gain adjustment (531) and a dynamic adjustment range of 530 – 600 nm using an optic filter module (FI 280 350, 1304A1) in a BMG PHERAstar® microplate reader. Subsequently, the substrate and enzyme solutions were mixed and incubated for 10 min before measuring the intrinsic Trp fluorescence again as described above. Enzyme-substrate (ES) complex formation was determined by subtracting the first enzyme only measurement from the measurement of the ES mixture and calculating the percentage of the quenching (change in intrinsic Trp fluorescence intensity (TFI)). Influences on fluorescence quenching not related to substrate binding were accounted for by using enzyme only controls for correction. The *K_d_* was estimated by plotting the corrected TFI values over the respective substrate concentration and computing a non-linear regression function through the data points.

#### *In silico* enzyme-substrate docking using HADDOCK2.4

Before uploading protein structures as PDB file to HADDOCK2.4, multiple occupancy side chains and water molecules were removed using the pdb_selaltloc PDB tool from https://wenmr.science.uu.nl/pdbtools/ and PyMOL Molecular Graphics System (Version 3.0 Schrödinger, LLC) respectively. The charge state of iron in PDB 6S7E and nickel in 7REI was adjusted to Fe^2+^ and Ni^2+^. The structures of enzyme substrates RAP2.12 and RGS4 were predicted using AlphaFold2 and truncated to 5- and 7-mer peptides (RAP2.12_2-6/8_ and RGS5_2-6_) using PyMOL. The Nt-Cys of the peptide substrate was renamed to CYF to ensure the sulfur atom of the Nt-Cys side chain can coordinate to the iron centre of the enzyme. Modified PDB files from enzymes and peptide substrates were uploaded to HADDOCK2.4. For the enzyme, residues D177 and Y183 (in *At*PCO4, D206 and Y212 in *Hs*ADO) and the metal co-factor were selected as active residues and “Remove buried active/passive residues from selection” was de-selected. For the peptide substrates, residue 2 was selected as active residue. The N-terminal amino group of the peptide was modified to -NH ^+^. The default option was used for all other settings.

### *At*PCO4 *in planta* experiments

#### Growth conditions

Col-0 was used as wildtype ecotype (seeds obtained from Dept. of Biology, University of Oxford). Plants were grown with a 16 hours photoperiod at 22 °C in a controlled environment room. At the end of a plant’s life cycle, it was transferred into a drying room until seed harvest. Plants used for the submergence experiment and RT-qPCR analysis were grown in heavier compost consisting of 1:1 modular seed and John Innes No.2 with ¼ fine vermiculite.

#### Preparation of plasmids for transformation into Arabidopsis plants

The mutations, C173A and Y183F, and the insertion of E135 to modify *At*PCO4-1 to *At*PCO4-2 were introduced into the entry vector pENTR223:PCO4 vector (Arabidopsis biological resource centre (ABRC) stock code G82286) that encodes *At*PCO4-1 using site-directed mutagenesis (described above, Table S1). After successful mutagenesis and amplification of the plasmids, the *AtPCO4-2* mutants were transferred into the destination vector pH7WG-promPCO4 (previously described in White *et al.* 2020) using Gateway Cloning, in particular, LR recombination. In more detail, entry and destination vector were incubated with LR clonase II in 1 x TE buffer (10 mM Tris-Cl, 1 mM EDTA, pH 8.0) for 16 – 18 hours at 25 °C. Proteinase K was added and the solution was incubated for 10 min at 37 °C to stop the reaction. The recombination solution was transformed into ccdB susceptible *E. coli* DH5α cells. Successfully recombined plasmids, i.e., pHWG-promPCO4-*At*PCO4-2-C173A and -Y183F, were confirmed by sequencing with primer sets attB1+2 and PCO4_F+R (Table S2) and transformed into the *Agrobacterium* strain GV3101. Successful colonies were confirmed by PCR using primer sets as described above.

#### Transformation of Arabidopsis plants

To create *4pco AtPCO4-2* WT and mutant plants, the floral dip method^64^ was used on the previously described triple knockout Arabidopsis plants^52^ in Col-0 background, *3pco+AtPCO5+/-* (*AtPCO1-/-, AtPCO2-/-, AtPCO4-/-* and *AtPCO5+/-*). This line had been used previously to generate *4pco* plants as *4pco* plants cannot self-fertilize^15,52^ and is based on T-DNA mutant lines from the Nottingham Arabidopsis Stock Centre (uNASC) as follows T-DNA insertion line N451210 for *At*PCO1 (At5g15120), N116554 for *At*PCO2 (At5g39890), N471015 for *At*PCO4 (At2g42670) and N684949 for *At*PCO5 (At3g58670).^52^ For the transformation, agrobacteria colonies containing the respective recombined pHWG-promPCO4-AtPCO4-2 plasmid were cultured in LB medium supplemented with 20 ng.mL^-1^ rifampicin, 50 ng.mL^-1^ gentamycin, 100 ng.mL^-1^ spectinomycin, and 300 ng.mL^-1^ streptomycin, at 28 °C and 140 – 160 rpm until the suspension turned turbid (∼18 – 24 hours). The cells were pelleted and resuspended in 5% sucrose supplemented with Silwett L-77. The inflorescences of *3pco+AtPCO5+/-* Arabidopsis plants were dipped into the agrobacterium-sucrose suspension and stored in the dark for 12 hours. This procedure was repeated after 7 days for a higher success rate. Once the growth cycle was finished, the seeds were harvested and dried.

#### Selection of 4pco AtPCO4-2 lines

After stratification of the seeds for 7 – 10 days, the seeds were spread on Hygromycin selection plates. Promising seedlings were transferred onto soil (3:1 modular seed compost and fine vermiculite). After 2 – 3 weeks the plants were genotyped for *AtPCO4* insertion as well as for *AtPCO5* WT and *AtPCO5* T-DNA using a piece of their leaf tissue (for primer list see Table S3) to determine *4pco AtPCO4-2* WT and mutant lines. The tissue was ground in DNA extraction buffer (200 mM Tris, 250 mM NaCl, 25 mM EDTA, 0.5% SDS, pH 7.5) and the suspension was centrifuged at maximal speed (∼15,000 x g) for 2 min. The DNA was precipitated from the supernatant by adding an equal amount of isopropanol and incubating the sample for 5 min. The DNA was pelleted at maximum speed (∼15,000 x g) for 5 min, the supernatant was discarded, and the pellet was dried. After resuspending the DNA in sterile ddH_2_O, the DNA was added to the PCR mix containing GoTaq mix, the appropriate primer pairs and DMSO. The PCR program included a 2 min denaturation at 95 °C, before 30 cycles of denaturation at 95 °C for 30 sec, annealing at 5 °C below the melting temperature (T_m_) of the primers for 30 sec and elongation at 72 °C for 1 min. The final step was a final elongation for 10 min at 72 °C before storing the PCR products at 4 °C until separating them on a 1% agarose gel. In addition, the sequence of the inserted *AtPCO4-2* was confirmed by sequencing (using primer pairs listed in Tables S2 and S3).

### RT-qPCR

RT-qPCR analysis was performed on copy DNA (cDNA) from 5.5-week-old plants. RNA was extracted from four plants per line using a Plant RNA purification kit. RNA concentrations were measured at an absorbance of 260 nm and calculated using a UV/Vis spectrophometer. The mRNA was transcribed into cDNA using a cDNA synthesis kit. The cDNA was mixed with the qPCRBIO SyGreen Mix Hi-ROX. Appropriate primer pairs (Table S4) were added and the reactions were run on a real-time PCR system. The cycle threshold (C_T_) of each target was normalized to the C_T_ of the housekeeping gene, UBQ10, to determine relative quantities of the transcript levels (ΔC_T_). The fold change (ΔΔC_T_) between mRNA present in *4pco AtPCO4-2* lines compared to Col-0 was calculated by normalizing the ΔC_T_ from the *4pco AtPCO4-2* lines to ΔC_T_ of Col-0.

### Submergence experiment

Plants were submerged in 26 cm tap water in black boxes that were additionally wrapped in aluminium foil to ensure complete darkness. Each box contained 10 x *4pco AtPCO4-2* WT or mutant plants and 10 x Col-0 plants as control. All plants were submerged at the 9-leaf stage at the end of a light cycle (after ∼15 hours of light). Dark controls were grown under aerobic conditions in the dark for the maximal submergence period. After de-submergence, the plants were covered with a transparent lid for 24 hours to enable recovery in humid conditions. A yes / no system was applied to score survival and recovery rates during the 10 days after submergence. On day 10 of recovery, the fresh weight of the rosettes was determined. The dry weight was measured after drying the rosettes at 50 °C until no weight difference between days could be observed.

## Acknowledgements

We gratefully acknowledge the help of Dr Dona Gunawardana (University of Oxford) for preparing chemically competent *E. coli* BL21 (DE3) cells and Dr Phil Holdship (University of Oxford) for his help with ICP-MS experiments. Plasmid pRK793 (addgene plasmid #8827) was a gift from Dr David Waugh^62^.

This work was supported by the European Research Council (European Union’s Horizon 2020 Research and Innovation Programme) Grant 864888 PCOMOD (A.D. and E.F.) and Grant 101001320 SYNOXYS (V.S. and F.L.). The work was also supported by the Biotechnology and Biological Sciences Research Council (UKRI-BBSRC) grant number BB/T008784/1 (S.H. and M.P.), grant number BB/Z516946/1 (F.L. and V.S.) and grant numbers BB/Y512953/1 and BB/X001059/1 (F.L.). M.P. was also supported by the Clarendon Scholarship at St Edmund Hall College.

## Author Contributions

A.D. and E.F. conceived and designed the study. A.D., A.d.G., S.H. and L.S.W. conducted experiments. M.P. generated *4pco AtPCO4-2* WT plants. V.S., F.L. and E.F. supervised the study. A.D., S.H. and E.F. analysed data. A.D. and A.d.G. prepared figures. A.D. and E.F. wrote the manuscript with input from all authors.

## Supplemental information

Document S1, containing

Figures S1 – S19, Tables S1 – S6 and Supplementary Discussion.

## References

1. Ahmed, F. et al. Waterlogging tolerance of crops: Breeding, mechanism of tolerance, molecular approaches, and future prospects. Biomed Res. Int. 2013, (2013). doi:10.1155/2013/963525.

2. Renziehausen, T., Dirr, A., Schmidt-Schippers, R., Flashman, E. & Schippers, J. Oxygen sensing and plant adaptation to flooding in a changing climate. Philos. Trans. R. Soc. B Biol. Sci. 380, (2025). doi:10.1098/rstb.2024.0238.

3. The Impact of Disasters and Crises on Agriculture and Food Security: 2021. The impact of disasters and crises on agriculture and food security: 2021 (FAO, 2021). doi:10.4060/cb3673en.

4. Voesenek, L. A. C. J. & Bailey-Serres, J. Flood adaptive traits and processes: an overview. New Phytol. 206, 57–73 (2015). doi:10.1111/nph.13209.

5. Sasidharan, R. et al. Signal Dynamics and Interactions during Flooding Stress. Plant Physiol. 176, 1106–1117 (2018). doi:10.1104/pp.17.01232.

6. Hartman, S. et al. Ethylene-mediated nitric oxide depletion pre-adapts plants to hypoxia stress. Nat. Commun. 10, 4020 (2019). doi:10.1038/s41467-019-12045-4.

7. Gibbs, D. J. et al. Homeostatic response to hypoxia is regulated by the N-end rule pathway in plants. Nature 479, 415–418 (2011). doi:10.1038/nature10534.

8. Licausi, F. et al. Oxygen sensing in plants is mediated by an N-end rule pathway for protein destabilization. Nature 479, 419–422 (2011). doi:10.1038/nature10536.

9. Weits, D. A. et al. Plant cysteine oxidases control the oxygen-dependent branch of the N-end-rule pathway. Nat. Commun. 5, 3425 (2014). doi:10.1038/ncomms4425.

10. Gibbs, D. J. et al. Oxygen-dependent proteolysis regulates the stability of angiosperm polycomb repressive complex 2 subunit VERNALIZATION 2. Nat. Commun. 9, 5438 (2018). doi:10.1038/s41467-018-07875-7.

11. Weits, D. A. et al. An apical hypoxic niche sets the pace of shoot meristem activity. Nature 569, 714–717 (2019). doi:10.1038/s41586-019-1203-6.

12. White, M. D. et al. Plant cysteine oxidases are dioxygenases that directly enable arginyl transferase-catalysed arginylation of N-end rule targets. Nat. Commun. 8, 14690 (2017). doi:10.1038/ncomms14690.

13. Dissmeyer, N. Conditional Protein Function via N-Degron Pathway–Mediated Proteostasis in Stress Physiology. Annu. Rev. Plant Biol. 70, 83–117 (2019). doi:10.1146/annurev-arplant-050718-095937.

14. Kim, L. et al. Structural analyses of the plant PRT6-UBR box in the Cys-Arg/N-degron pathway and insights into the plant submergence response. *bioRxiv* at (2022). doi:10.1101/2022.08.19.504472.

15. White, M. D. et al. Structures of Arabidopsis thaliana oxygen-sensing plant cysteine oxidases 4 and 5 enable targeted manipulation of their activity. Proc. Natl. Acad. Sci. 117, 23140–23147 (2020). doi:10.1073/pnas.2000206117.

16. Weits, D. A. et al. Acquisition of hypoxia inducibility by oxygen sensing N-terminal cysteine oxidase in spermatophytes. Plant. Cell Environ. 46, 322–338 (2023). doi:10.1111/pce.14440.

17. Yeung, E., Bailey-Serres, J. & Sasidharan, R. After The Deluge: Plant Revival Post-Flooding. Trends Plant Sci. 24, 443–454 (2019). doi:10.1016/j.tplants.2019.02.007.

18. Schmidt, R. R. et al. Low-oxygen response is triggered by an ATP-dependent shift in oleoyl-CoA in *Arabidopsis*. Proc. Natl. Acad. Sci. 115, E12101–E12110 (2018). doi:10.1073/pnas.1809429115.

19. Fan, B. et al. Calcium-dependent activation of CPK12 facilitates its cytoplasm-to-nucleus translocation to potentiate plant hypoxia sensing by phosphorylating ERF-VII transcription factors. Mol. Plant 16, 979–998 (2023). doi:10.1016/j.molp.2023.04.002.

20. Hsiao, P.-Y., Zeng, C.-Y. & Shih, M.-C. Group VII ethylene response factors forming distinct regulatory loops mediate submergence responses. Plant Physiol. 194, 1745– 1763 (2024). doi:10.1093/plphys/kiad547.

21. Gibbs, D. J. et al. Nitric Oxide Sensing in Plants Is Mediated by Proteolytic Control of Group VII ERF Transcription Factors. Mol. Cell 53, 369–379 (2014). doi:10.1016/j.molcel.2013.12.020.

22. Ramon, M. et al. Default Activation and Nuclear Translocation of the Plant Cellular Energy Sensor SnRK1 Regulate Metabolic Stress Responses and Development. Plant Cell 31, 1614–1632 (2019). doi:10.1105/tpc.18.00500.

23. Cho, H., Lu, M. J. & Shih, M. The SnRK1-eIFiso4G1 signaling relay regulates the translation of specific mRNAs in Arabidopsis under submergence. New Phytol. 222, 366–381 (2019). doi:10.1111/nph.15589.

24. Kunkowska, A. B. et al. Target of rapamycin signaling couples energy to oxygen sensing to modulate hypoxic gene expression in *Arabidopsis*. Proc. Natl. Acad. Sci. 120, e2212474120 (2023). doi:10.1073/pnas.2212474120.

25. Yeung, E. et al. A stress recovery signaling network for enhanced flooding tolerance in *Arabidopsis thaliana*. Proc. Natl. Acad. Sci. 115, E6085–E6094 (2018). doi:10.1073/pnas.1803841115.

26. Pospíšil, P. Production of reactive oxygen species by photosystem II. Biochim. Biophys. Acta - Bioenerg. 1787, 1151–1160 (2009). doi:10.1016/j.bbabio.2009.05.005.

27. Fukao, T., Yeung, E. & Bailey-Serres, J. The Submergence Tolerance Regulator SUB1A Mediates Crosstalk between Submergence and Drought Tolerance in Rice. Plant Cell 23, 412–427 (2011). doi:10.1105/tpc.110.080325.

28. Tamang, B. G., Magliozzi, J. O., Maroof, M. A. S. & Fukao, T. Physiological and transcriptomic characterization of submergence and reoxygenation responses in soybean seedlings. Plant. Cell Environ. 37, 2350–2365 (2014). doi:10.1111/pce.12277.

29. Luo, F.-L. et al. De-submergence responses of antioxidative defense systems in two wetland plants having escape and quiescence strategies. J. Plant Physiol. 169, 1680– 1689 (2012). doi:10.1016/j.jplph.2012.06.015.

30. Setter, T. L., Bhekasut, P. & Greenway, H. Desiccation of leaves after de-submergence is one cause for intolerance to complete submergence of the rice cultivar IR 42. Funct. Plant Biol. 37, 1096 (2010). doi:10.1071/FP10025.

31. Yuan, L. et al. Multi-stress resilience in plants recovering from submergence. Plant Biotechnol. J. 21, 466–481 (2023). doi:10.1111/pbi.13944.

32. Alpuerto, J. B., Hussain, R. M. F. & Fukao, T. The key regulator of submergence tolerance, SUB1A, promotes photosynthetic and metabolic recovery from submergence damage in rice leaves. Plant. Cell Environ. 39, 672–684 (2016). doi:10.1111/pce.12661.

33. Locke, A. M., Barding, G. A., Sathnur, S., Larive, C. K. & Bailey-Serres, J. Rice SUB1A constrains remodelling of the transcriptome and metabolome during submergence to facilitate post-submergence recovery. Plant. Cell Environ. 41, 721–736 (2018). doi:10.1111/pce.13094.

34. Lin, C. C. et al. Regulatory cascade involving transcriptional and N-end rule pathways in rice under submergence. Proc. Natl. Acad. Sci. U. S. A. 116, 3300–3309 (2019). doi:10.1073/pnas.1818507116.

35. Singh, S., Mackill, D. J. & Ismail, A. M. Responses of *SUB1* rice introgression lines to submergence in the field: Yield and grain quality. F. Crop. Res. 113, 12–23 (2009). doi:10.1016/j.fcr.2009.04.003.

36. Singh, U. S. et al. Field performance, dissemination, impact and tracking of submergence tolerant (Sub1) rice varieties in South Asia. SABRAO J. Breed. Genet. 45, 112–131 (2013).

37. Mendiondo, G. M. et al. Enhanced waterlogging tolerance in barley by manipulation of expression of the N-end rule pathway E3 ligase PROTEOLYSIS6. Plant Biotechnol. J. 14, 40–50 (2016). doi:10.1111/pbi.12334.

38. Yu, F. et al. A group VII ethylene response factor gene, ZmEREB180, coordinates waterlogging tolerance in maize seedlings. Plant Biotechnol. J. 17, 2286–2298 (2019). doi:10.1111/pbi.13140.

39. Qi, H. et al. ZmEREB180 modulates waterlogging tolerance in maize by regulating root development and antioxidant gene expression. Plant Biotechnol. J. 23, 2062–2064 (2025). doi:10.1111/pbi.70030.

40. Taylor-Kearney, L. J. & Flashman, E. Targeting plant cysteine oxidase activity for improved submergence tolerance. Plant J. 109, 779–788 (2022). doi:10.1111/tpj.15605.

41. Chen, Z. et al. Molecular basis for cysteine oxidation by plant cysteine oxidases from *Arabidopsis thaliana*. J. Struct. Biol. 213, 107663 (2021). doi:10.1016/j.jsb.2020.107663.

42. Gunawardana, D. M., Heathcote, K. C. & Flashman, E. Emerging roles for thiol dioxygenases as oxygen sensors. FEBS J. 289, 5426–5439 (2022). doi:10.1111/febs.16147.

43. Jiramongkol, Y. et al. An mRNA-display derived cyclic peptide scaffold reveals the substrate binding interactions of an N-terminal cysteine oxidase. Nat. Commun. 16, 4761 (2025). doi:10.1038/s41467-025-59960-3.

44. Ye, S. et al. An Insight into the Mechanism of Human Cysteine Dioxygenase. J. Biol. Chem. 282, 3391–3402 (2007). doi:10.1074/jbc.M609337200.

45. Wang, Y., Shin, I., Li, J. & Liu, A. Crystal structure of human cysteamine dioxygenase provides a structural rationale for its function as an oxygen sensor. J. Biol. Chem. 297, 101176 (2021). doi:10.1016/j.jbc.2021.101176.

46. White, M. D., Kamps, J. J. A. G., East, S., Taylor Kearney, L. J. & Flashman, E. The plant cysteine oxidases from *Arabidopsis thaliana* are kinetically tailored to act as oxygen sensors. J. Biol. Chem. 293, 11786–11795 (2018). doi:10.1074/jbc.RA118.003496.

47. Dirr, A., Gunawardana, D. M. & Flashman, E. Kinetic Measurements to Investigate the Oxygen-Sensing Properties of Plant Cysteine Oxidases. in Oxygen Sensing: Methods and Protocols (ed. Weinert, E. E.) 207–230 (Springer US, New York, NY, 2023). doi:10.1007/978-1-0716-3080-8_13.

48. Schofield, C. J. & Ratcliffe, P. J. Oxygen sensing by HIF hydroxylases. Nat. Rev. Mol. Cell Biol. 5, 343–354 (2004). doi:10.1038/nrm1366.

49. Clifton, I. J. et al. Structural studies on 2-oxoglutarate oxygenases and related double-stranded β-helix fold proteins. J. Inorg. Biochem. 100, 644–669 (2006). doi:10.1016/j.jinorgbio.2006.01.024.

50. de Visser, S. P. & Straganz, G. D. Why Do Cysteine Dioxygenase Enzymes Contain a 3-His Ligand Motif Rather than a 2His/1Asp Motif Like Most Nonheme Dioxygenases? J. Phys. Chem. A 113, 1835–1846 (2009). doi:10.1021/jp809700f.

51. Fernandez, R. L. et al. The Crystal Structure of Cysteamine Dioxygenase Reveals the Origin of the Large Substrate Scope of This Vital Mammalian Enzyme. Biochemistry 60, 3728–3737 (2021). doi:10.1021/acs.biochem.1c00463.

52. Masson, N. et al. Conserved N-terminal cysteine dioxygenases transduce responses to hypoxia in animals and plants. Science 365, 65–69 (2019). doi:10.1126/science.aaw0112.

53. Gunawardana, D. M., Southern, D. A. & Flashman, E. Measuring plant cysteine oxidase interactions with substrates using intrinsic tryptophan fluorescence. Sci. Rep. 14, 1–10 (2024). doi:10.1038/s41598-024-83508-y.

54. Bui, L. T. et al. Differential submergence tolerance between juvenile and adult Arabidopsis plants involves the ANAC017 transcription factor. Plant J. 104, 979–994 (2020). doi:10.1111/tpj.14975.

55. Liu, K. et al. Silver lining to a climate crisis in multiple prospects for alleviating crop waterlogging under future climates. Nat. Commun. 14, 765 (2023). doi:10.1038/s41467-023-36129-4.

56. Paul, M. V. et al. Oxygen Sensing via the Ethylene Response Transcription Factor RAP2.12 Affects Plant Metabolism and Performance under Both Normoxia and Hypoxia. Plant Physiol. 172, 141–153 (2016). doi:10.1104/pp.16.00460.

57. Kourelis, J. et al. Bioengineering secreted proteases converts divergent Rcr3 orthologs and paralogs into extracellular immune co-receptors. Plant Cell 36, 3260–3276 (2024). doi:10.1093/plcell/koae183.

58. Schuster, M. et al. Enhanced late blight resistance by engineering an EpiC2B-insensitive immune protease. Plant Biotechnol. J. 22, 284–286 (2024). doi:10.1111/pbi.14209.

59. Vicente, J. et al. The Cys-Arg/N-End Rule Pathway Is a General Sensor of Abiotic Stress in Flowering Plants. Curr. Biol. 27, 3183–3190.e4 (2017). doi:10.1016/j.cub.2017.09.006.

60. Wang, K., Guo, H. & Yin, Y. AP2/ERF transcription factors and their functions in *Arabidopsis* responses to abiotic stresses. Environ. Exp. Bot. 222, 105763 (2024). doi:10.1016/j.envexpbot.2024.105763.

61. Zheng, L. An efficient one-step site-directed and site-saturation mutagenesis protocol. Nucleic Acids Res. 32, e115 (2004). doi:10.1093/nar/gnh110.

62. Kapust, R. B. et al. Tobacco etch virus protease: mechanism of autolysis and rational design of stable mutants with wild-type catalytic proficiency. Protein Eng. Des. Sel. 14, 993–1000 (2001). doi:10.1093/protein/14.12.993.

63. Taylor-Kearney, L. J. et al. Plant Cysteine Oxidase Oxygen-Sensing Function Is Conserved in Early Land Plants and Algae. ACS Bio Med Chem Au 2, 521–528 (2022). doi:10.1021/acsbiomedchemau.2c00032.

64. Clough, S. J. & Bent, A. F. Floral dip: a simplified method for *Agrobacterium*-mediated transformation of *Arabidopsis thaliana*. Plant J. 16, 735–743 (1998). doi:10.1046/j.1365-313x.1998.00343.x.

