## Supplementary material for "Engineering reduced activity in the oxygen-sensing *Arabidopsis thaliana* plant cysteine oxidase 4 enzyme results in improved flood resilience": Document S1

**SUPPLEMENTARY INFORMATION**


**Authors/affiliations:**

Anna Dirr[1], Arianna Del Greco[2,3], Simon Howe[2], Leah S. Walker[2], Monica Perri[2], Vinay Shukla[2], Francesco Licausi[2], Emily Flashman[2*]

[1]Department of Chemistry, University of Oxford, United Kingdom

[2]Department of Biology, University of Oxford, United Kingdom

[3]University of Pisa, Biology Department, Via L. Ghini 13 - 56126, Pisa, Italy

### Supplementary Figures

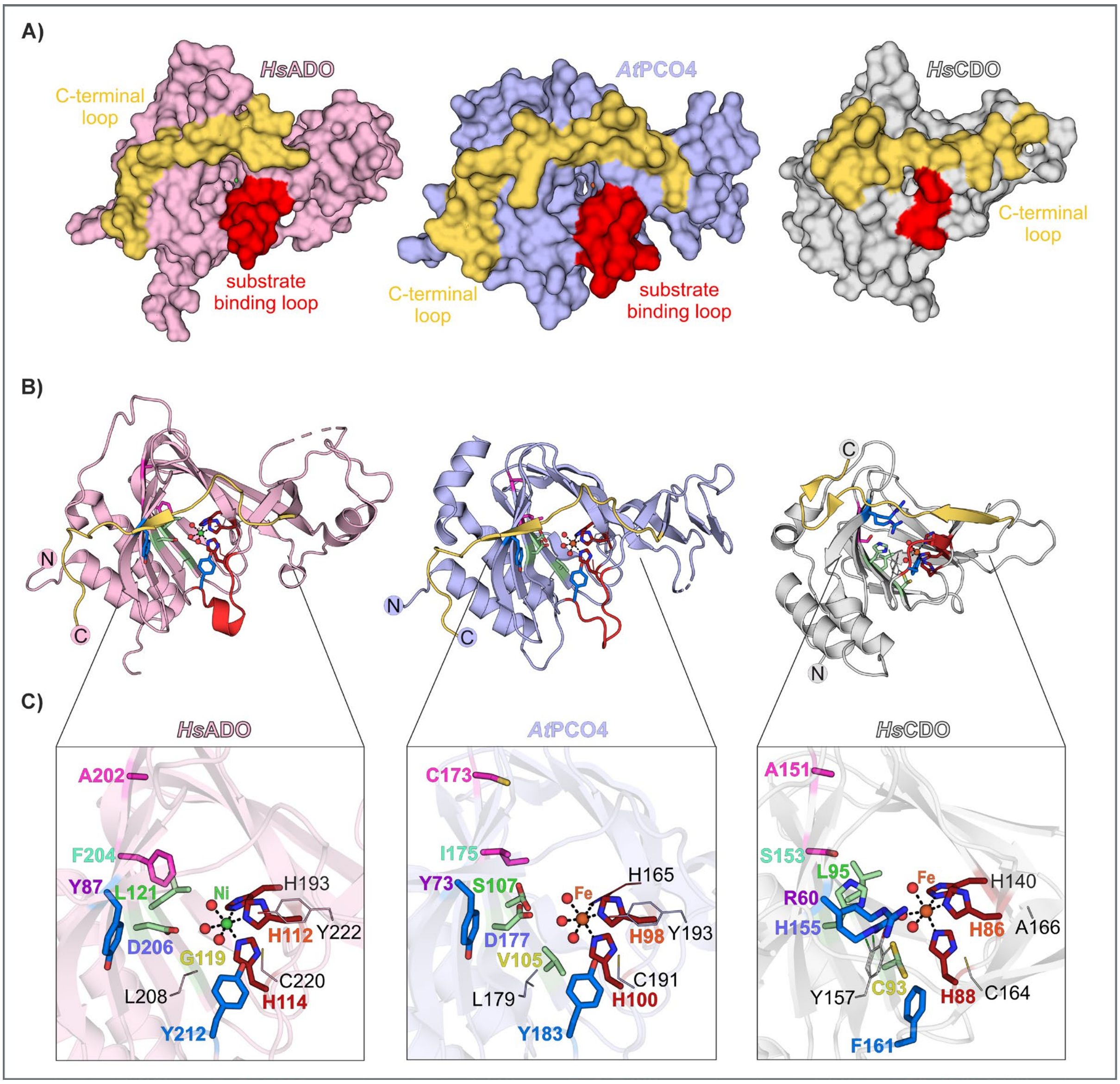


**Figure S1. Structural comparison of *At*PCO4 with the thiol dioxygenases *Hs*ADO and *Hs*CDO-1.**
**(A)** Surface and **(B)** cartoon presentation of X-ray crystal structures from *Hs*ADO (PDB: 7REI), *At*PCO4 (PDB: 6S7E, structure shown is *At*PCO4-1 but residues numbered according to *At*PCO4-2 to align with main text) and *Hs*CDO-1 (PDB: 6CDH). **(C)** Cartoon representation of the active site of each enzyme with residues combined according to their location (same colour), i.e., red: iron coordinating His-triad, blue: residues at potential substrate entry site, pink: residues *trans* to substrate entry site, green: residues *trans* to His-triad.

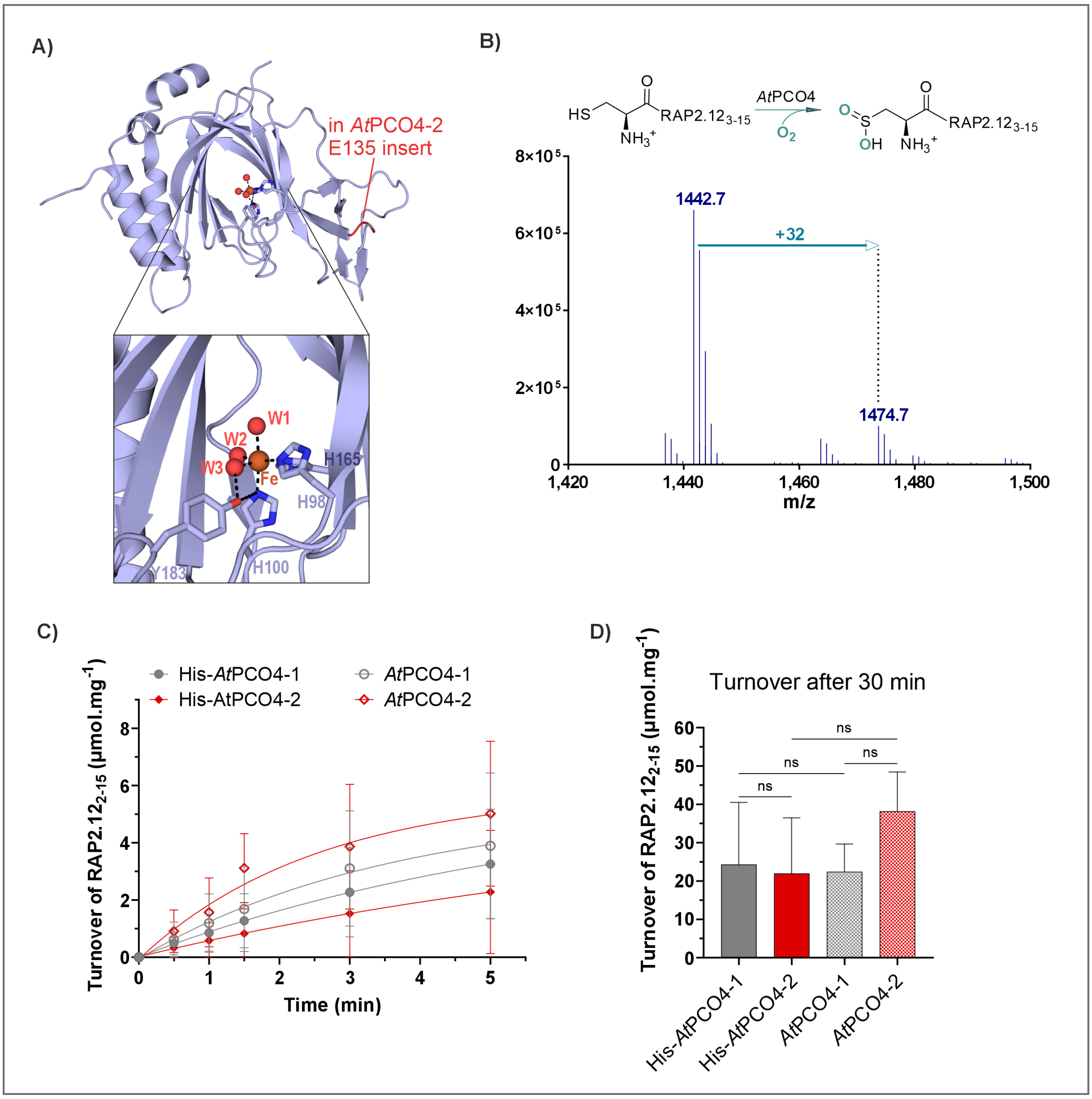


**Figure S2. Structural difference between *At*PCO4-1 and -2 and their *in vitro* activities.**
**(A)** Cartoon presentation of *At*PCO4-1 with indicated additional residue in isoform 2 (PDB: 6S7E). **(B)** Deconvoluted mass spectrum of the dioxygenated (+ 32 Da) RAP2.12$_{2\text{-}15}$ peptide (1442.7 Da) catalysed by *At*PCO4-2 (10 min reaction time). **(C)** and **(D)** Activity comparison of *At*PCO4-1 and -2 with and without His-tag. Activities are shown as turnover of µmol substrate per mg enzyme. The assays were performed in technical triplicates (n = 3) with 0.4 µM enzyme from two purified batches of *At*PCO4-1 and one batch of *At*PCO4-2 at 25 °C in reaction buffer (50 mM bis-tris propane (pH 8), 50 mM NaCl) with 5 µM $FeSO_4$, 1 mM L-ascorbate, 1 mM TCEP and 200 µM RAP2.12$_{2\text{–}15}$. His-*At*PCO4-1 and -2 converted 63% and 43% substrate (~ turnover number of 157 and 109, respectively). *At*PCO4-1 and -2 converted 69% and 76% substrate (~ turnover number of 172 and 190, respectively). The data showed that differences between the enzymes were not significant (ns) for $P < 0.05$ after performing one-way ANOVA with Holm-Šídák's test (GraphPad Prism, Version 10.1.2).

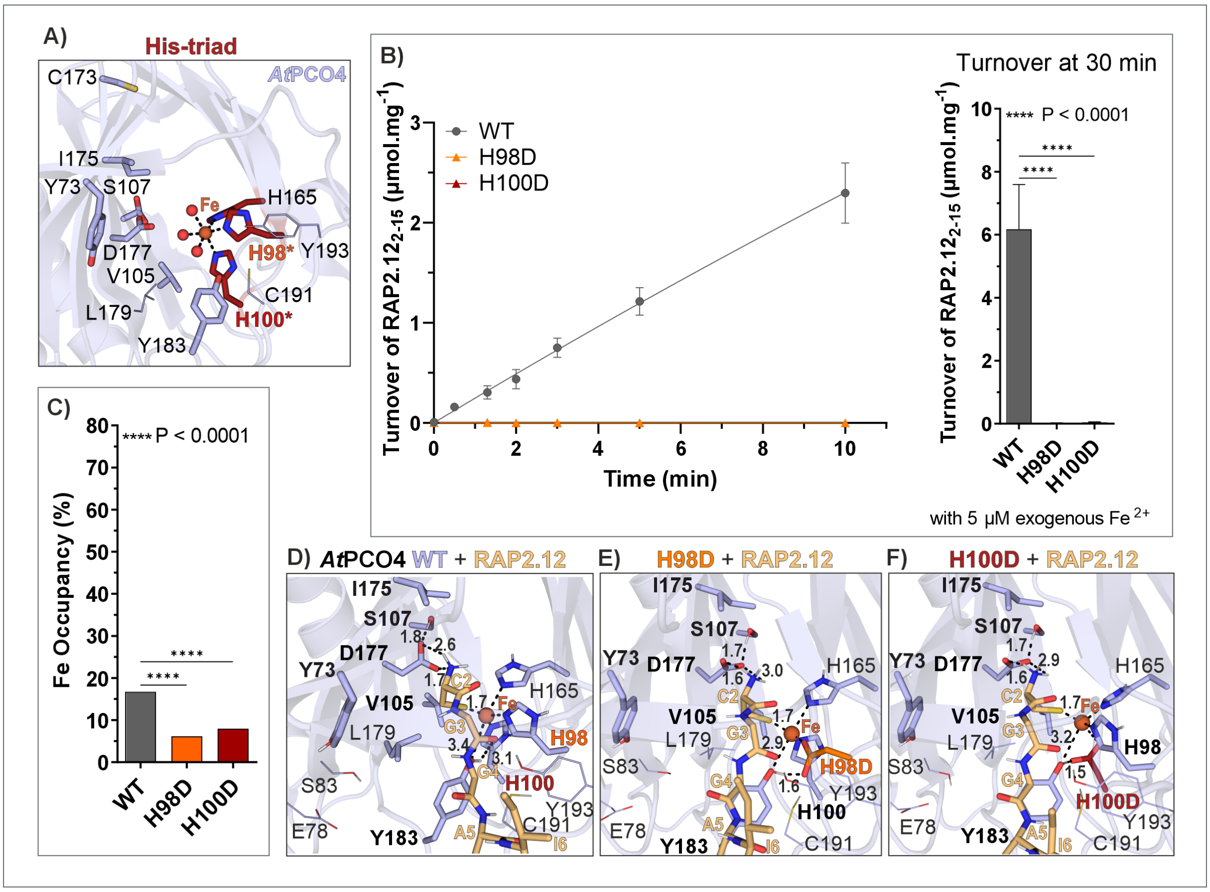


**Figure S3. Iron occupancy, activity and substrate binding models of His-triad variants H98D and H100D.**

**(A)** Cartoon representation of *At*PCO4 active site obtained from a crystal structure (PDB: 6S7E) with His-triad residues highlighted in red. **(B)** *At*PCO4 WT (n = 9) and variant (n = 3) activity with 200 µM RAP2.12$_{2\text{-}15}$ in assay buffer (50 mM bis-tris-propane, pH 8, 50 mM NaCl) with 1 mM TCEP and 1 mM L-ascorbate. Statistical significance was determined using one-way ANOVA and Holm-Šídák's testing to correct for multiple comparisons (GraphPad Prism). **(C)** Iron occupancy determined using ICP-MS for WT and variant H100D (n = 2) and a colorimetric assay for variant H98D (n = 3). **(D)** – **(F)** *In silico* docking results of *At*PCO4 WT and His-triad variants (based on PDB 6S7E and created using the mutagenesis tool in PyMOL) with RAP2.12$_{2\text{-}6}$ using HADDOCK2.4. The predicted models are presented as cartoon and other active site residues investigated are indicated as sticks and with bold letters.

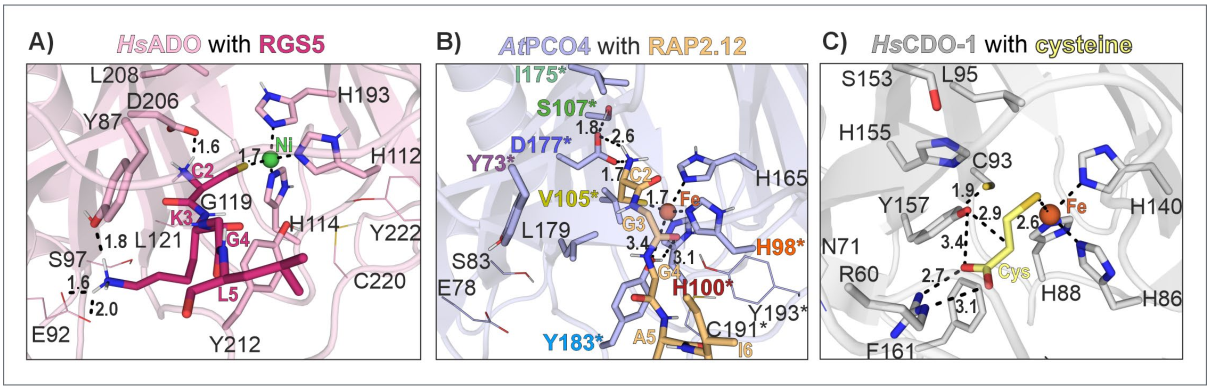


**Figure S4. Substrate binding of *Hs*ADO, *At*PCO4-2 and *Hs*CDO-1.**
Cartoon representation of the active site of **(A)** *Hs*ADO (PDB: 7REI) with its 4-mer peptide substrate RGS5$_{2–5}$ and **(B)** *At*PCO4-2 (PDB: 6S7E) with its 5-mer peptide substrate RAP2.12$_{2–6}$ docked using HADDOCK 2.4.[1] **(C)** Cartoon representation of the active site of *Hs*CDO-1 with its substrate cysteine (PDB: 6BGF) positioned in the crystal structure. Key residues and substrates are represented as sticks. Active site metals are represented as green (Ni) or orange (Fe) spheres. Dashed lines represent distances between atoms (Å).

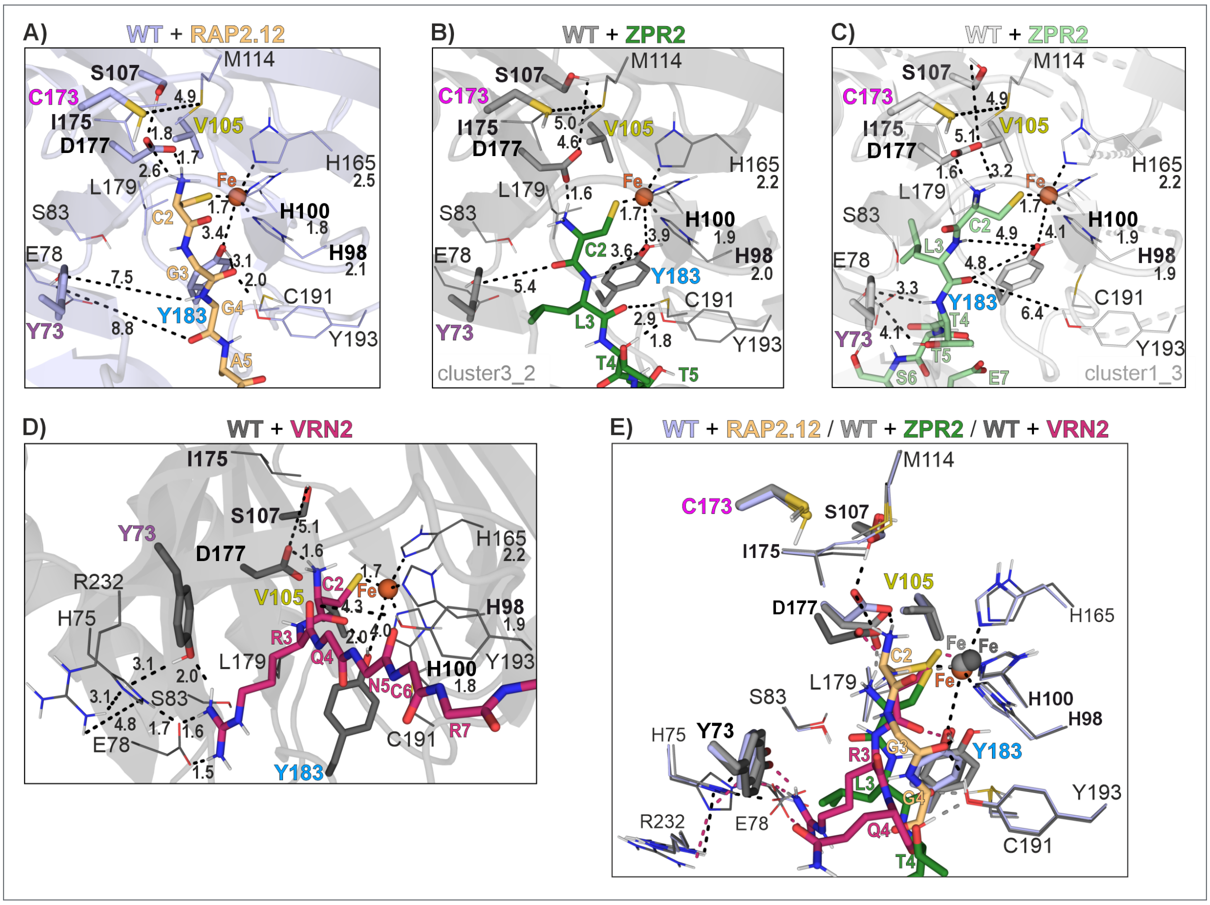


**Figure S5. *In silico* docking models of ERFVII and non-ERFVII substrates to *At*PCO4-2 WT using Haddock2.1.**

Cartoon representation of the active site of *At*PCO4-2 (PDB: 6S7E) WT with its 5-mer peptide substrates **(A)** RAP2.12$_{2–6}$, **(E)** ZPR2$_{2\text{-}6}$ and **(D)** VRN2$_{2\text{-}7}$ docked using HADDOCK 2.4.[1] **(B)** and **(C)** Two different clusters of the docked ZPR2$_{2\text{-}6}$ substrate. **(E)** Overlay of the docked RAP2.12$_{2–6}$, ZPR2$_{2\text{-}6}$ and VRN2$_{2\text{-}7}$ substrates in the active site of *At*PCO4-2 WT. ZPR2$_{2\text{-}6}$ and VRN2$_{2\text{-}7}$ are peptides representing the Cys2-initiating N-terminal sequences of known PCO substrates Little Zipper 2 and Vernalisation 2.[2–5]

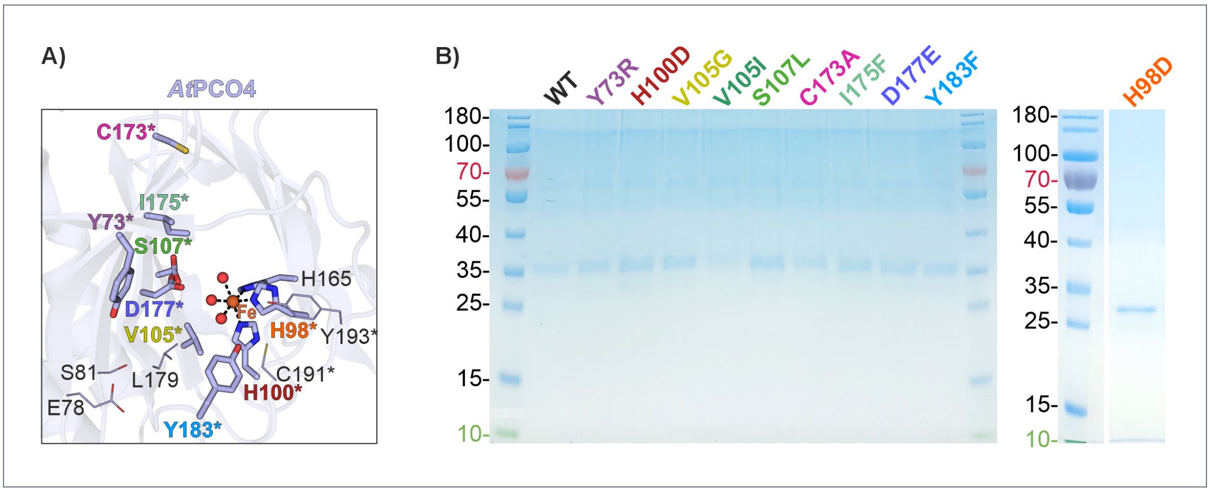


**Figure S6. Overview of recombinant purified *At*PCO4 WT and variants.**
**(A)** Cartoon representation of the *At*PCO4 active site (PDB: 6S7E). **(B)** Left, SDS-PAGE gel of *At*PCO4 WT and active site variants after storage at - 80 °C and, on the right, of a SEC fraction from *At*PCO4 H98D stained with QuickBlue Protein Stain.

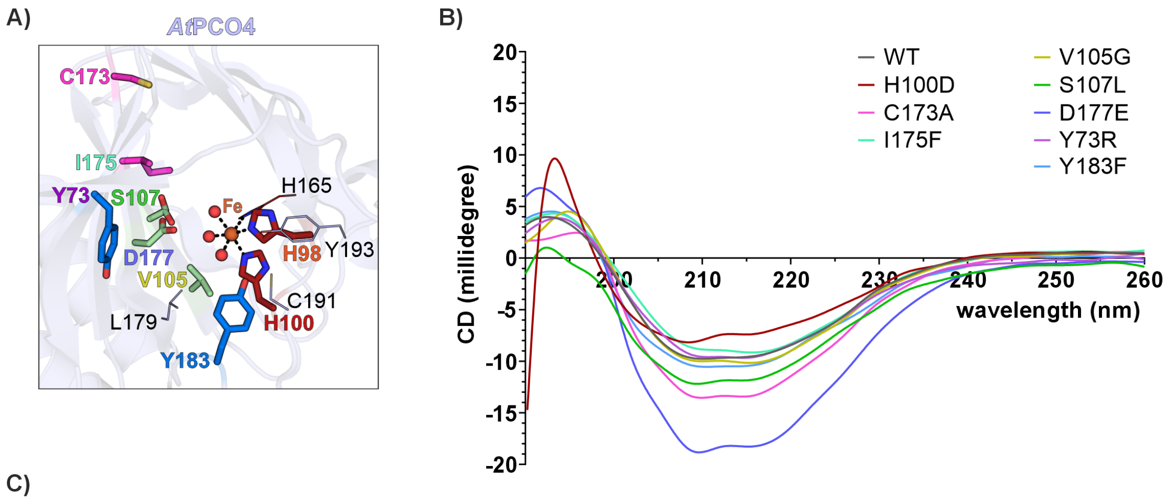


C)

| DichroWeb results using K2D method | | | | |
|---|---|---|---|---|
| | α-helix | β-sheet | random coil | |
| WT | 5% | 47% | 48% | |
| WT for H100D | 5% | 47% | 48% | |
| H100D | 5% | 47% | 48% | residue coordinating the iron centre |
| C173A | 5% | 47% | 48% | residues *trans* to substrate entry site |
| I175F | 5% | 47% | 48% | |
| V105G | 5% | 47% | 48% | residues *trans* to His-triad |
| S107L | 5% | 47% | 48% | |
| D177E | 5% | 47% | 48% | |
| Y73R | 5% | 47% | 48% | residues at substrate entry site |
| Y183F | 5% | 47% | 48% | |

**Figure S7. CD spectra of active site variants.**
**(A)** Cartoon representation of the *At*PCO4 active site (PDB: 6S7E). **(B)** CD spectra of *At*PCO4 enzymes after subtracting the baseline. The CD spectra were measured at a protein concentration of 0.2 mg.mL$^{-1}$ in 10 mM $KH_2PO_4$ (pH 7.5), 260 nm monochromator wavelength, 1.0 nm bandwidth, 185 – 260 nm in 0.5 steps, 0.5 s time-per-point and at 25 °C. **(C)** Predictions of *At*PCO4 enzyme secondary structures from experimental CD results using the K2D method on DichroWeb[6]. Maximum error for all predictions was determined as > 0.22.

A)

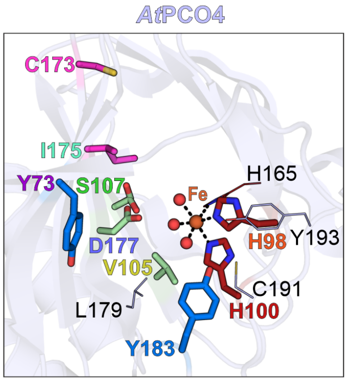


B)

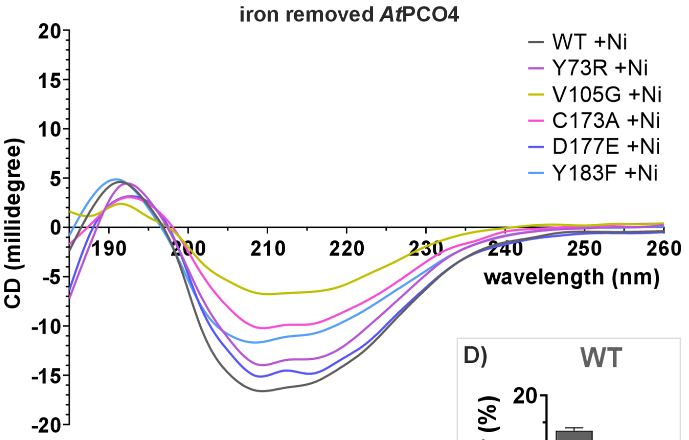


C)

| DichroWeb results using K2D method | | | | |
|---|---|---|---|---|
| | α-helix | β-sheet | random coil | |
| WT | 5% | 47% | 48% | |
| C173A | 5% | 47% | 48% | residue *trans* to substrate entry site |
| V105G | 5% | 47% | 48% | residues *trans* to His-triad |
| D177E | 5% | 47% | 48% | |
| Y73R | 5% | 47% | 48% | residues at substrate entry site |
| Y183F | 5% | 47% | 48% | |

D)

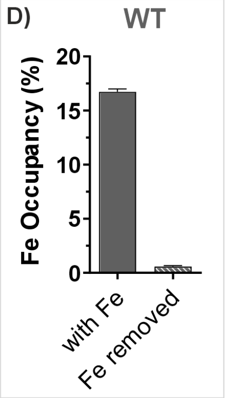


**Figure S8. CD spectra of iron removed AtPCO4 enzymes supplemented with nickel.**
**(A)** Cartoon representation of the *At*PCO4 active site (PDB: 6S7E). **(B)** CD spectra of iron removed and with nickel substituted (+Ni) *At*PCO4 enzymes after subtracting the baseline. The CD spectra were measured at a protein concentration of 0.2 mg.mL$^{-1}$ in 10 mM $KH_2PO_4$ (pH 7.5), 260 nm monochromator wavelength, 1.0 nm bandwidth, 185 – 260 nm in 0.5 steps, 0.5 s time-per-point and at 25 °C. **(C)** Predictions of *At*PCO4 enzyme secondary structures from experimental CD results using the K2D method on DichroWeb[6]. Maximum error for all predictions was determined as > 0.22. **(D)** Iron occupancy of *At*PCO4 WT after removing the iron during the purification process determined by ICP-MS (technical replicates, n = 2).

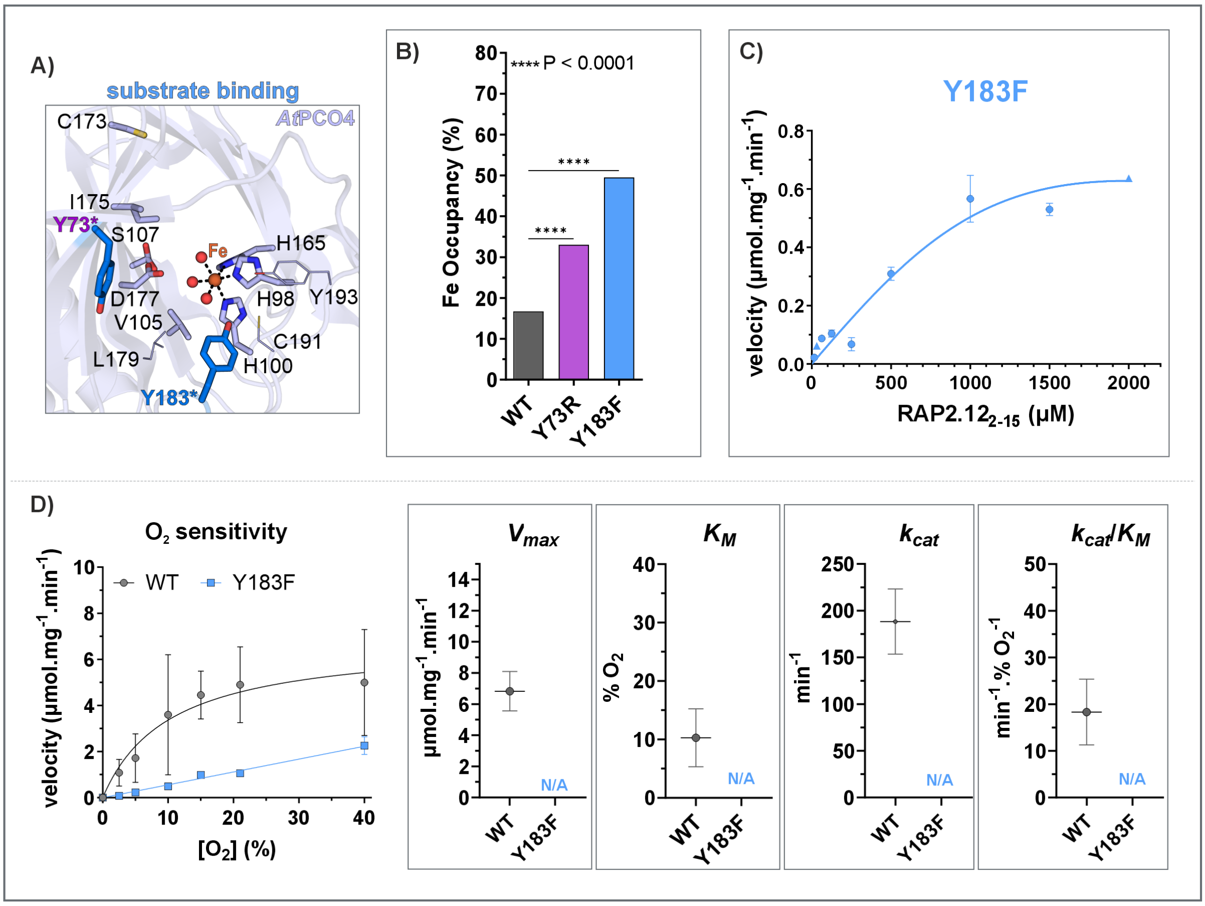


**Figure S9. Iron occupancy and $O_2$ sensitivity of variants with mutations at the substrate entry site.**

**(A)** Cartoon representation of *At*PCO4 active site obtained from a crystal structure (PDB: 6S7E) with substrate entry site residues highlighted in bright blue. **(B)** Iron occupancy determined using ICP-MS (n = 2). Statistical significance was determined using one-way ANOVA and Holm-Šídák's testing to correct for multiple comparisons (GraphPad Prism). **(C)** Initial velocities of *At*PCO4 Y183F (in technical triplicates, n = 3) from 0 – 2 min time courses plotted against $RAP2_{2\text{-}15}$ peptide substrate concentrations. Solid line is the fit of the Michaelis-Menten equation to the data. Standard deviation is indicated by error bars. **(D)** $O_2$ sensitivity of AtPCO4 WT and variant Y183F measured after 1 min reaction time (left, replicates: 2 x n = 3). Standard deviation is indicated by error bars. Michaelis-Menten constants were estimated using GraphPad Prism. Error bars of Michaelis-Menten constants indicate the standard error of the mean. Statistical significance could not be evaluated because the Michaelis-Menten model was not applicable (N/A) for variant Y183F.

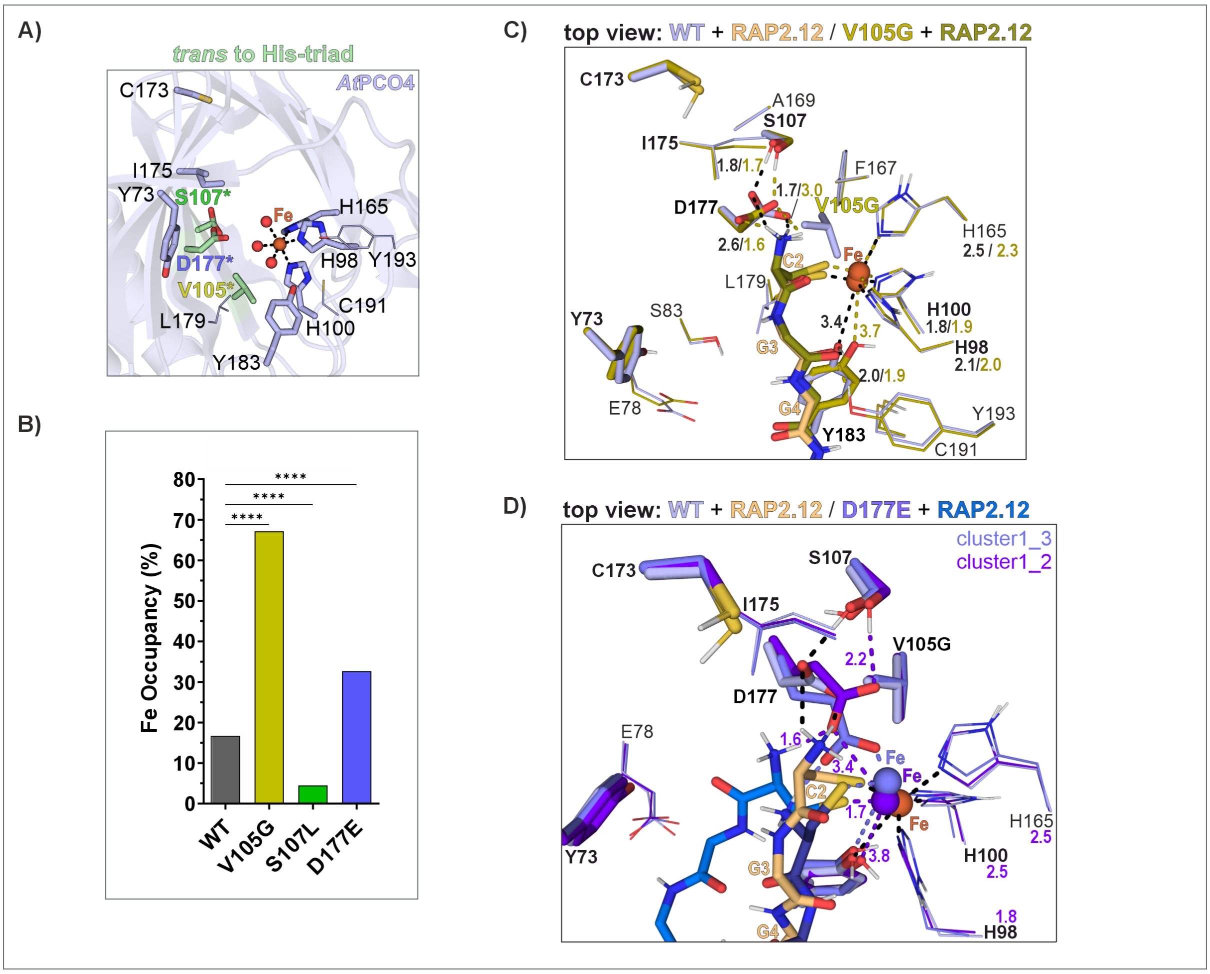


**Figure S10. Substrate binding and iron occupancy of variants with mutations *trans* to His-triad. (A)** Cartoon representation of *At*PCO4 active site obtained from a crystal structure (PDB: 6S7E). Residues that are positioned *trans* to the His-triad are highlighted in green. **(B)** Iron occupancy determined using ICP-MS (n = 2). Statistical significance was determined using one-way ANOVA and Holm-Šídák's testing to correct for multiple comparisons (****: P < 0.0001, GraphPad Prism). **(C)** and **(D)** *in silico* docking results of *At*PCO4 WT and variants (based on PDB 6S7E and created using the mutagenesis tool in PyMOL) with RAP2.12$_{2\text{-}8}$ using HADDOCK2.4.[1,7] The predicted variant models (in panel **(C)** for variant V105G, in panel **(D)** cluster1_3 and cluster1_2 for variant D177E) were aligned with the WT model and are presented as cartoon. Other investigated active site residues are indicated in bold letters. **(D)** The docking models for variant D177E indicate that E177 can either coordinate the iron (clusters1_3) or still coordinate the substrate (cluster1_2).

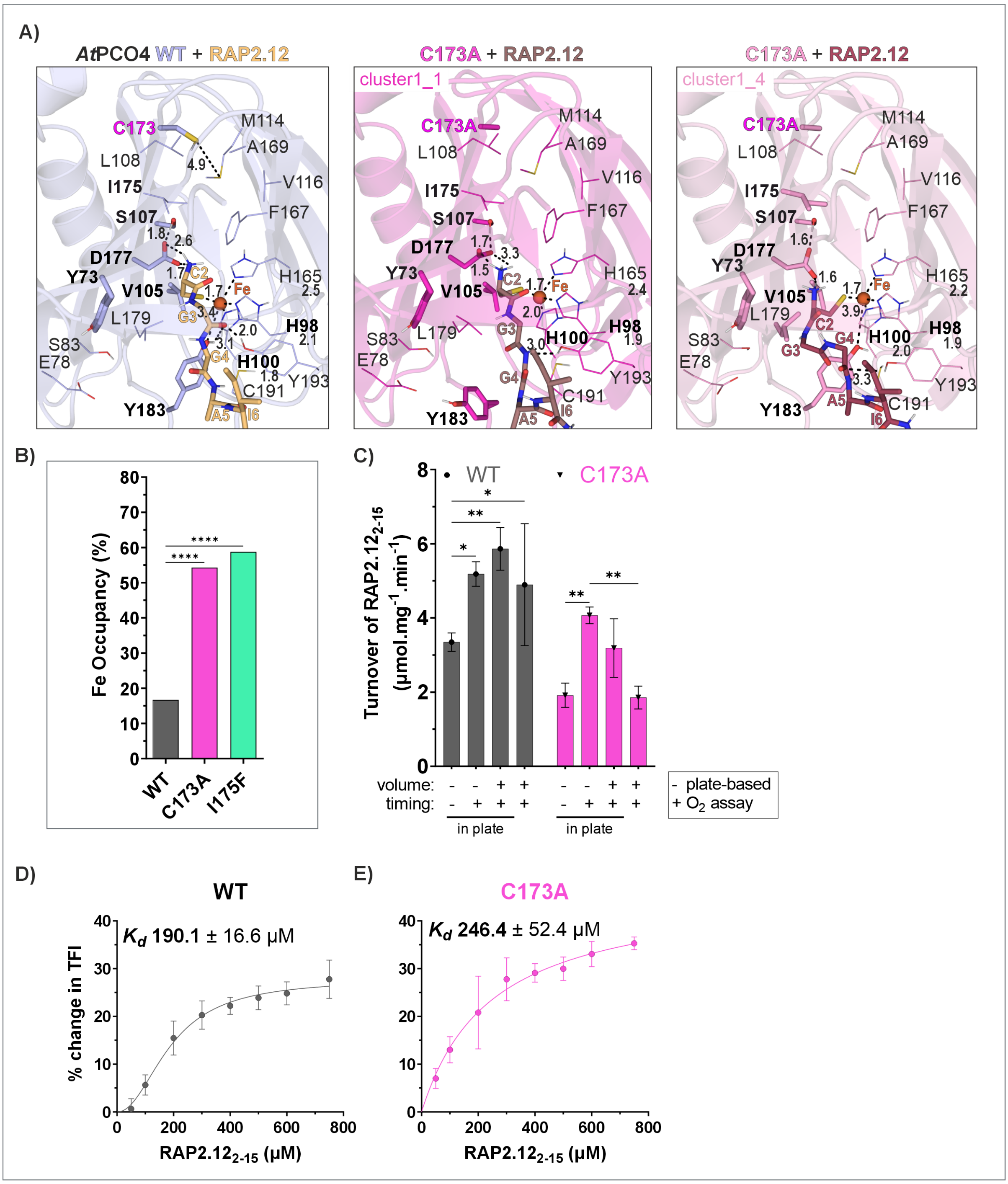


**Figure S11. Substrate binding and iron occupancy of variants with mutations *trans* to the substrate entry site.**

**(A)** Structures showing *in silico* docking results of *At*PCO4 WT and variant C173A (based on PDB 6S7E and created using the mutagenesis tool in PyMOL) with RAP2.12$_{2\text{-}8}$ predicted using HADDOCK2.4[1,7] and presented as cartoons. Other investigated active site residues are indicated as sticks and in bold letters. **(B)** Iron occupancy determined using ICP-MS (n = 2). Statistical significance was determined using one-way ANOVA and Holm-Šídák's testing to correct for multiple comparisons (****: $P < 0.0001$, GraphPad Prism). **(C)** Comparison of enzyme activity between plate-based assays and $O_2$ assays for which vials were used to enable measurements at specific $O_2$ concentrations. The activity of 0.1 µM *At*PCO4 WT and variant C173A (n = 3, n = 5 for $O_2$ assay) was measured with buffer, substrate and enzyme volumes and incubation timings used in the different assay setups at optimal reaction conditions (RAP2.12$_{2\text{–}15}$, iron, L-ascorbate) after 1 min in assay buffer (50 mM bis-tris-propane, pH 8, 50 mM NaCl) with 5 mM TCEP at 21% $O_2$. **(D)** and **(E)** dissociation curves of RAP2.12$_{2\text{-}15}$ from *At*PCO4 enzymes acquired using changes in intrinsic tryptophan fluorescence.

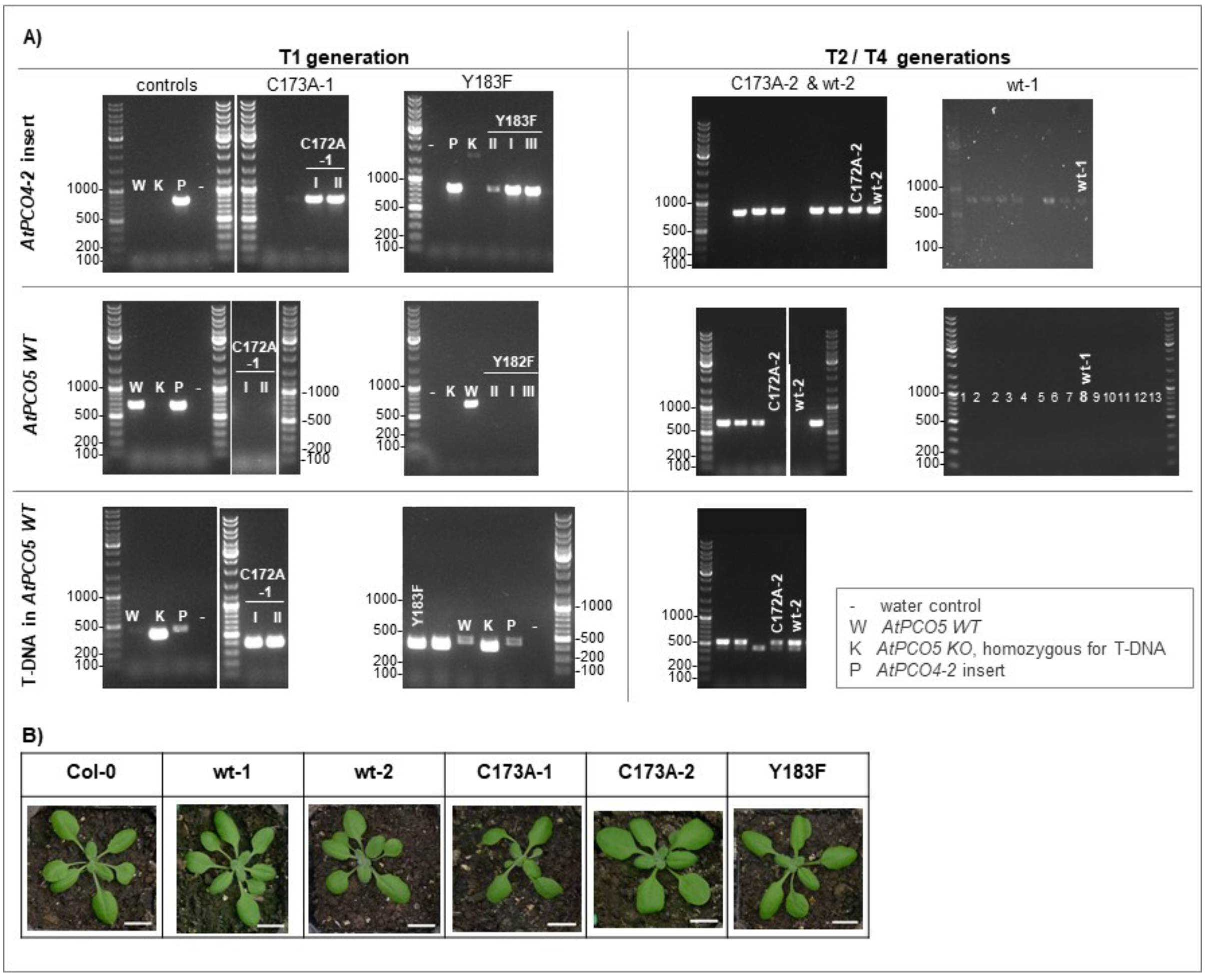


**Figure S12. Complementation of *4pco* Arabidopsis plants with *AtPCO4 WT*, *C173A* and *Y183F*.** **(A)** Genotyping results of *4pco* knockout plants complemented with *AtPCO4 WT* and mutants. Amplified PCR products that correspond to the *AtPCO4* insert are shown in the top panel. Plants homozygous for *AtPCO5*$^{-/-}$ were identified by an amplified PCR product for the T-DNA insert (bottom panel) and lack of amplification for *AtPCO5 WT* (middle panel). The labels indicate the plant line and roman letters indicate PCR results from repeated DNA extractions from the same plant. The generations on top specify in which generation the plant was homozygous for *AtPCO5*$^{-/-}$ after transforming *3pco+AtPCO5*$^{+/-}$ with *AtPCO4 WT* and mutants. The full names of the lines when identified as *4pco AtPCO4* WT or mutant were as follows: wt-1 = L22-17-19-8, wt-2 = 14-14-14-h6, C173A-1 = C2, C173A-2 = A23-h6, Y183F = A20. **(B)** Phenotypes of Col-0 and the final lines used for further analysis: *4pco AtPCO4 WT* lines (wt-1, wt-2), *AtPCO4-C173A* lines (C173A-1, C173A-2) and *AtPCO4-Y183F* line 3-weeks post-sowing. Scale bar: 1 cm.

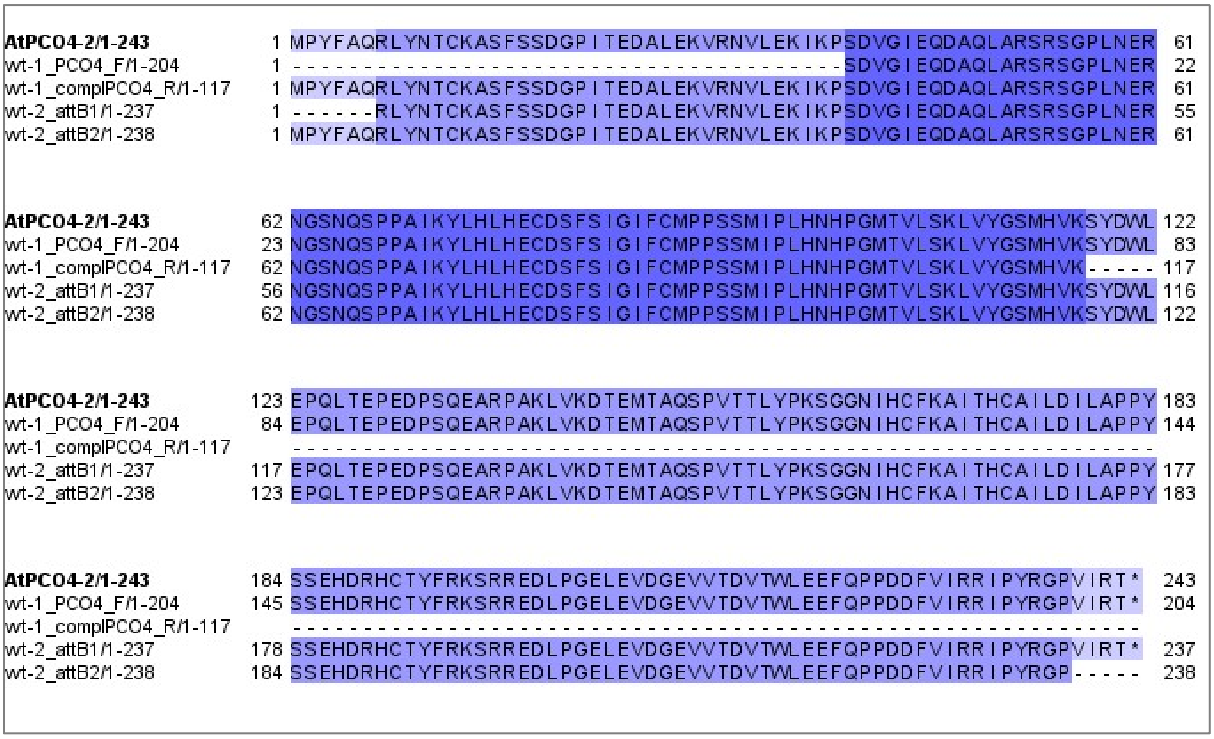


**Figure S13. Sequencing results from complemented lines wt-1 and -2.**
DNA sequences were translated using SnapGene Viewer (Version 6.0.7) and aligned using Clustal Omega[8]. Residues are coloured according to their percentage identity in Jalview 2 (Version 2.11.3.2). Primer pairs used for sequencing are indicated on the left.

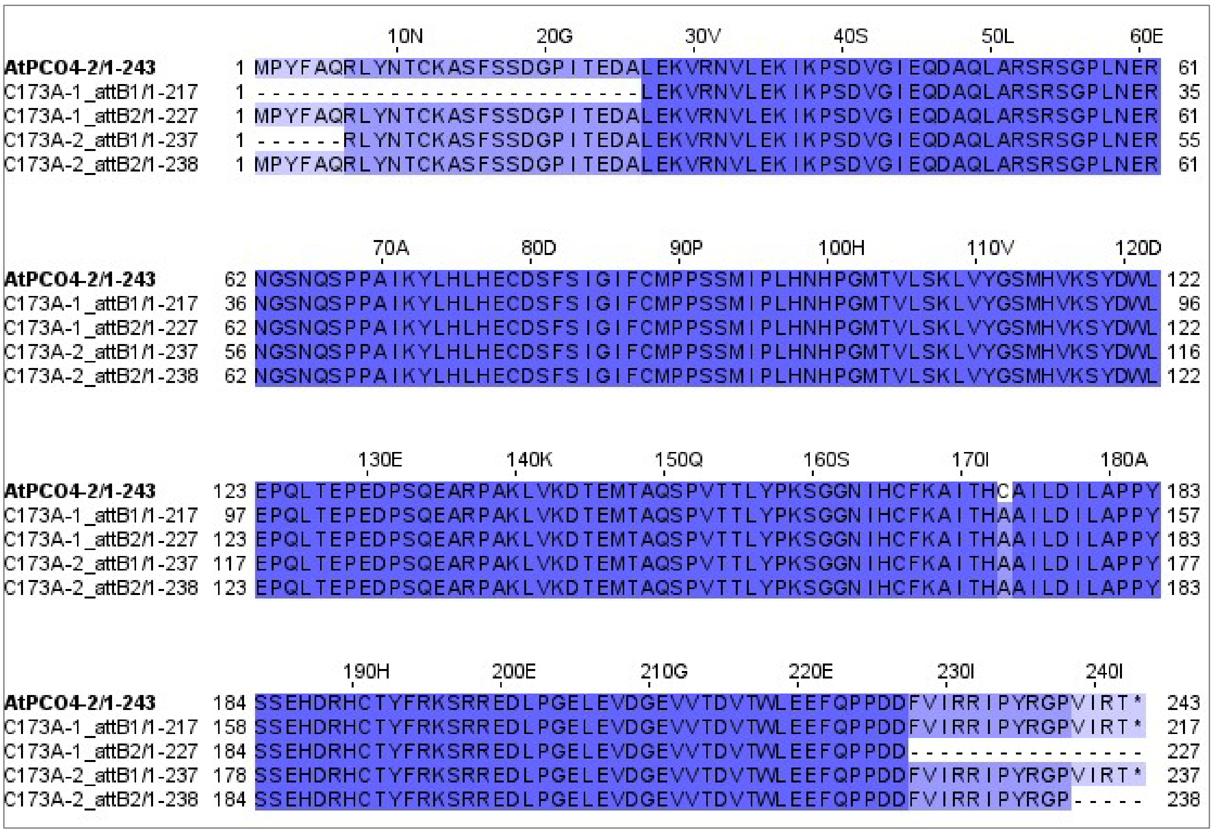


**Figure S14. Sequencing results from complemented lines C173A-1 and -2.**

DNA sequences were translated using SnapGene Viewer (Version 6.0.7) and aligned using Clustal Omega[8]. Residues are coloured according to their percentage identity in Jalview 2 (Version 2.11.3.2). Primer pairs used for sequencing are indicated on the left.

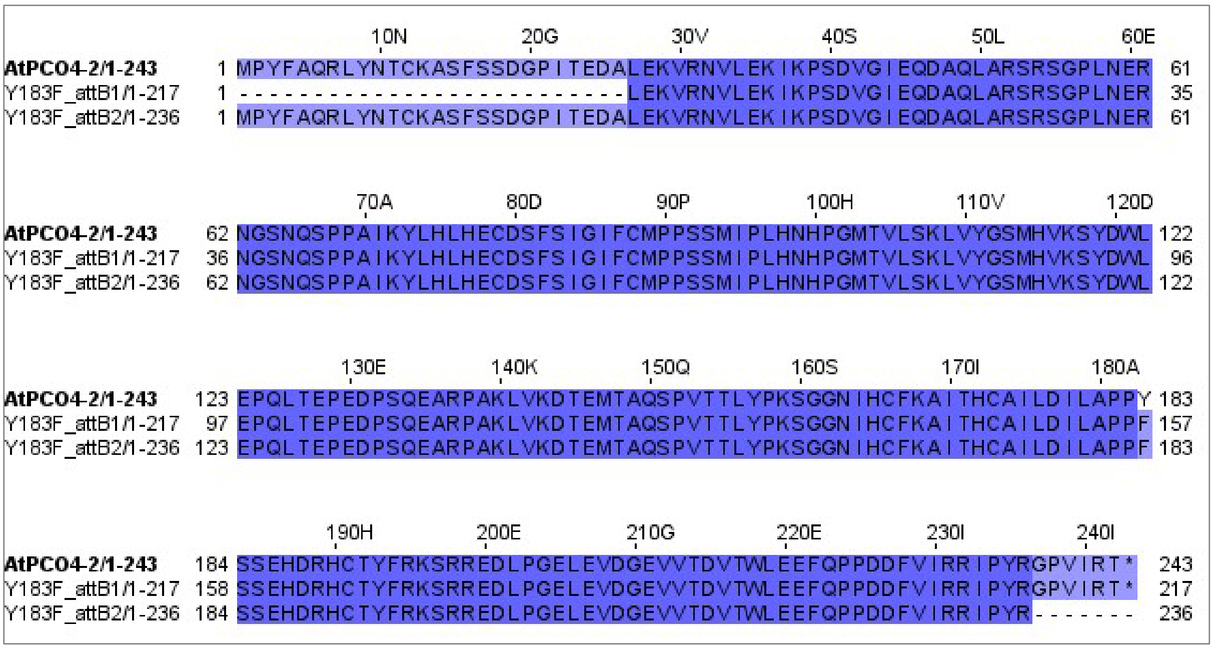


**Figure S15. Sequencing results from complemented line Y183F.**
DNA sequences were translated using SnapGene Viewer (Version 6.0.7) and aligned using Clustal Omega[8]. Residues are coloured according to their percentage identity in Jalview 2 (Version 2.11.3.2). Primer pairs used for sequencing are indicated on the left.

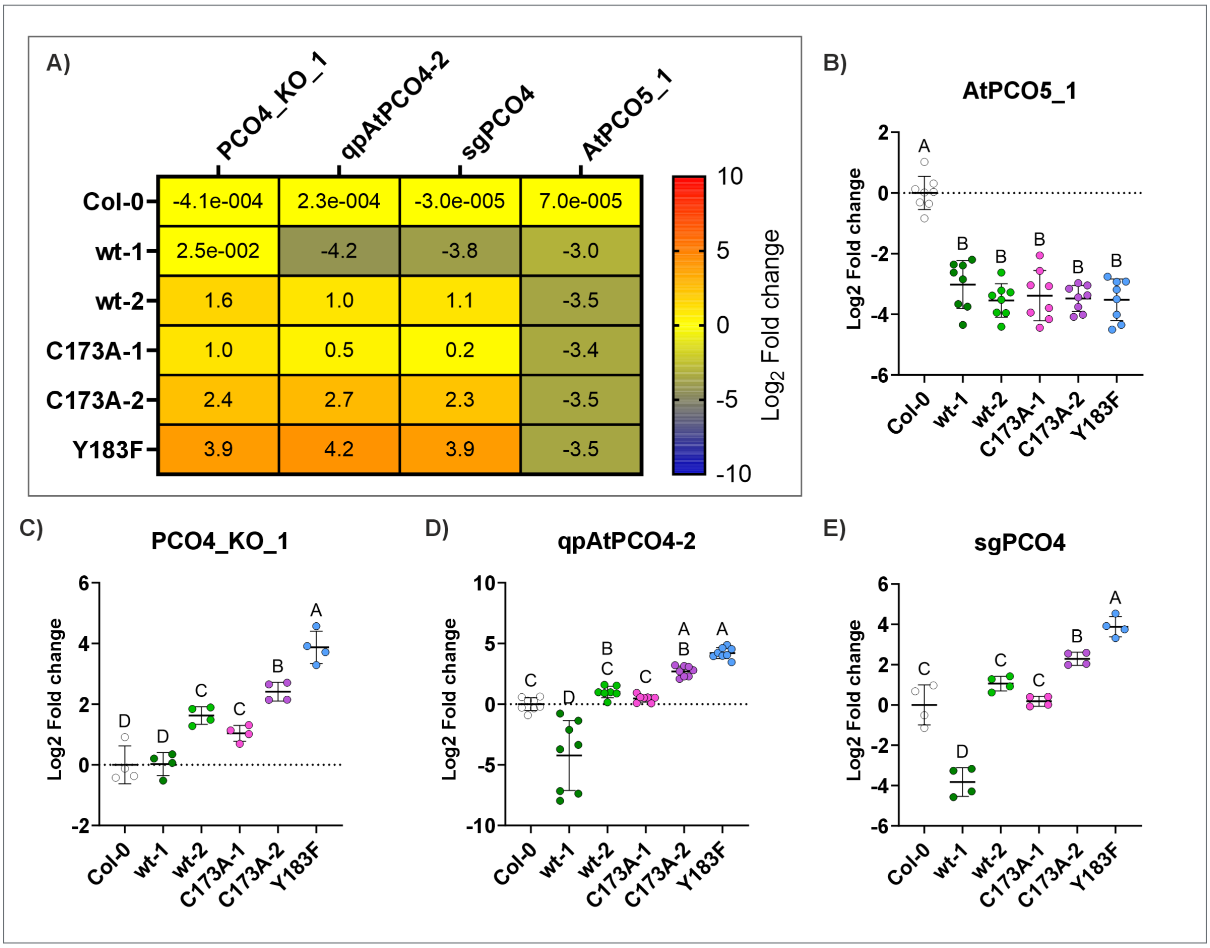


**Figure S16. *AtPCO4* and *5* transcript levels of 5.5-week-old *4pco AtPCO4* plants.**
RT-qPCR results were normalised to transcript levels of the housekeeping gene UBQ10 and Col-0. **(A)** Transcript levels of inserted *AtPCO4* were determined using three different primer sets: **(C)** PCO4_KO_1, **(D)** qpAtPCO4-2 (two independent experiments) and **(E)** sgPCO4. The latter two primer sets covered a similar region +/- 2 bp. Transformed *AtPCO4* transcript levels were similar between all primer sets except for line wt-1 for which either no transcript levels (PCO4_KO_1) similar to Col-0 or reduced levels were determined (qpAtPCO4-2, sgPCO4). **(B)** *AtPCO5* transcript levels were measured using primer set AtPCO5_1 to confirm *pco5*$^{-/-}$ and therefore the *4pco* genotype (two independent experiments). **(B)** – **(E)** Statistical significance ($P < 0.05$) was obtained using one-way ANOVA and the Holm-Šídák's test for multiple comparison testing (GraphPad Prism).

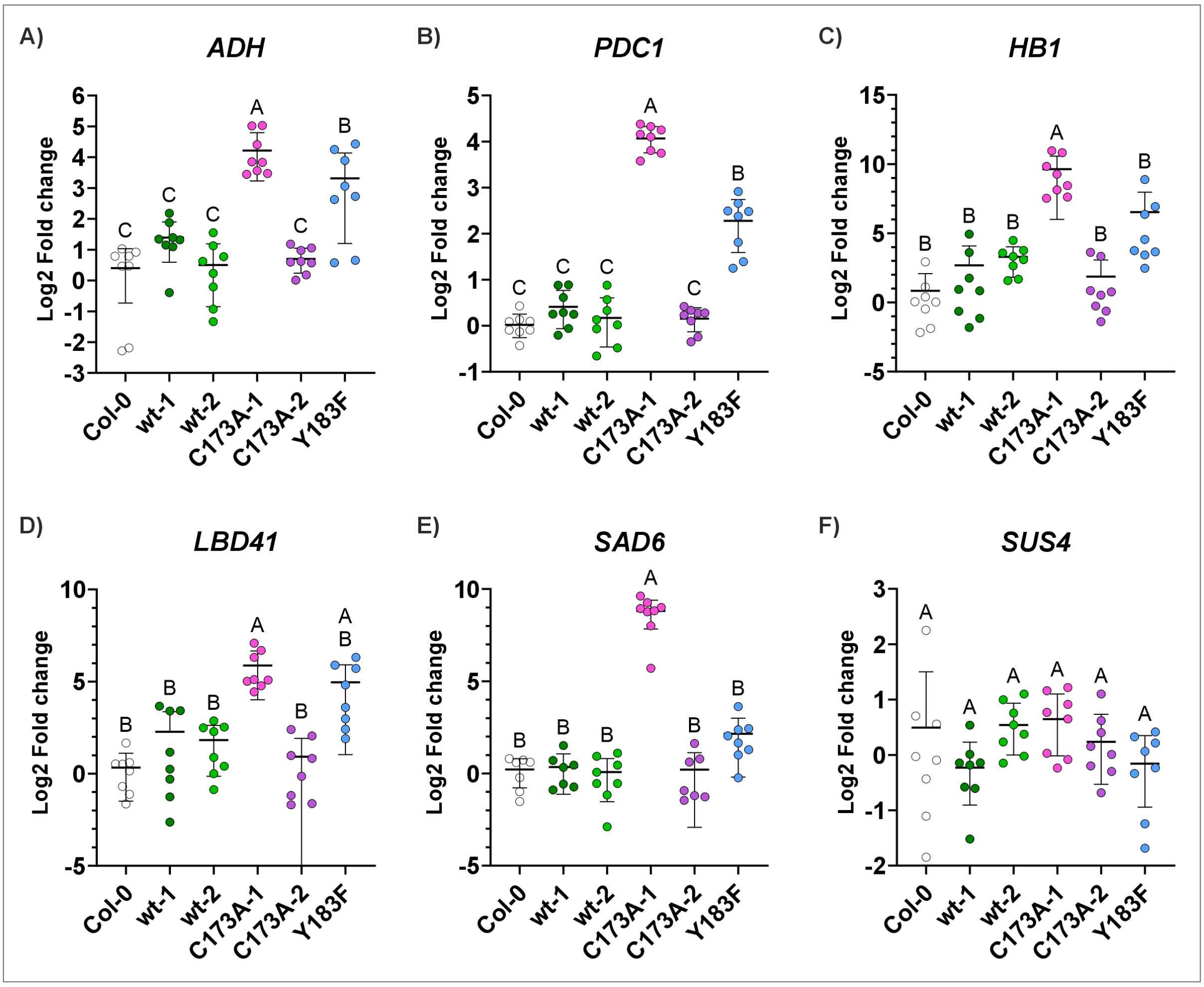


**Figure S17. Statistical significance of differences in HRG transcript levels of *4pco AtPCO4* plants.**

Change in transcript levels of HRGs in 5.5-week-old *4pco AtPCO4* and Col-0 plants. Two independent experiments with two technical replicates of four biological samples each (n = 2 x (2 x 4)). Statistical significance ($P < 0.05$) was obtained using one-way ANOVA and the Holm-Šídák's test for multiple comparison testing (GraphPad Prism).

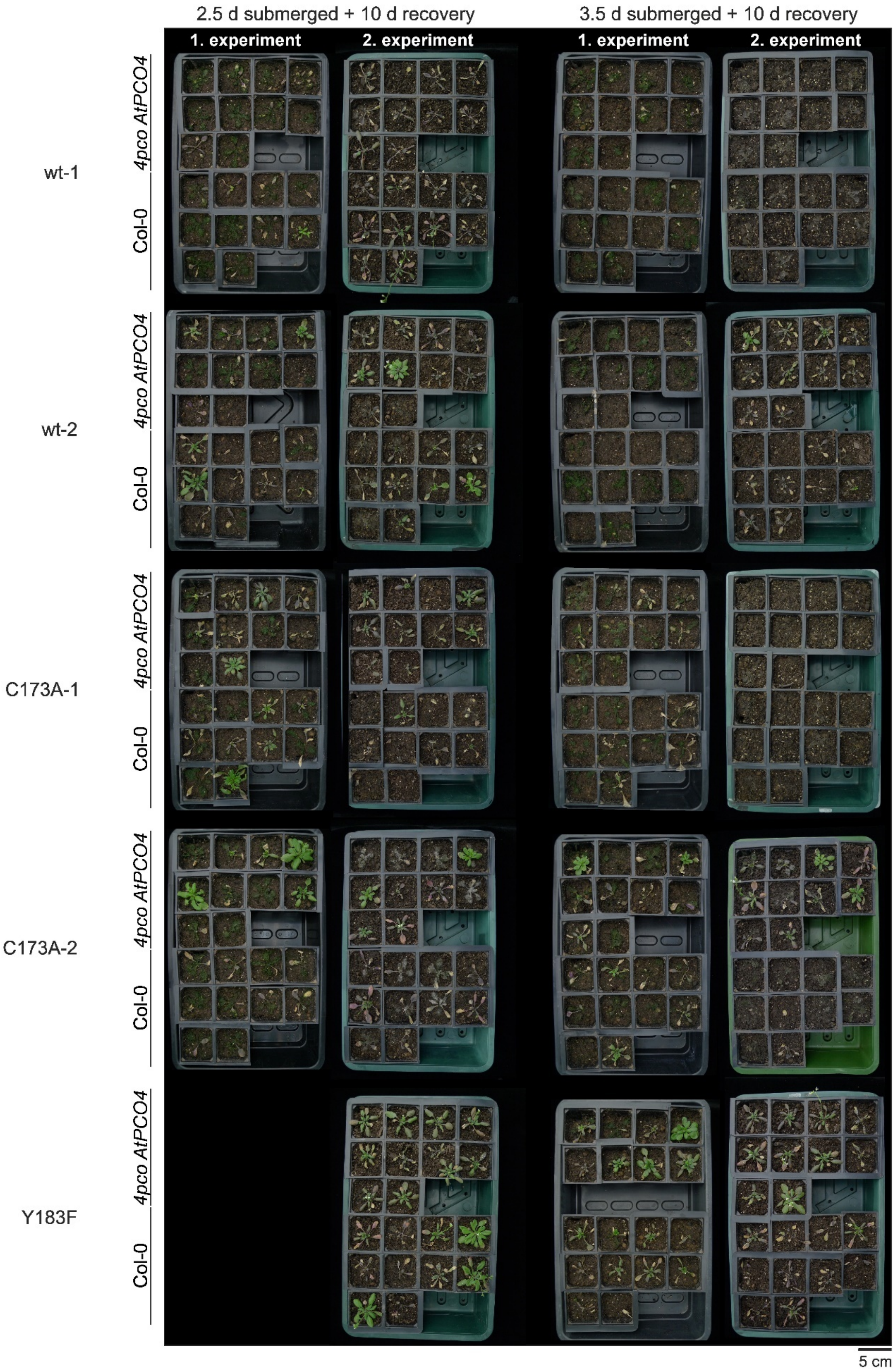


**Figure S18. Submergence recovery of all *4pco* lines and their Col-0 control plants.**
Photos of all submerged plants after 10 days of recovery. Each tray contains a complemented *4pco* line and Col-0 as control. For line Y183F, only eight plants were available for submergence in the first experiment which were submerged for 3.5 days.

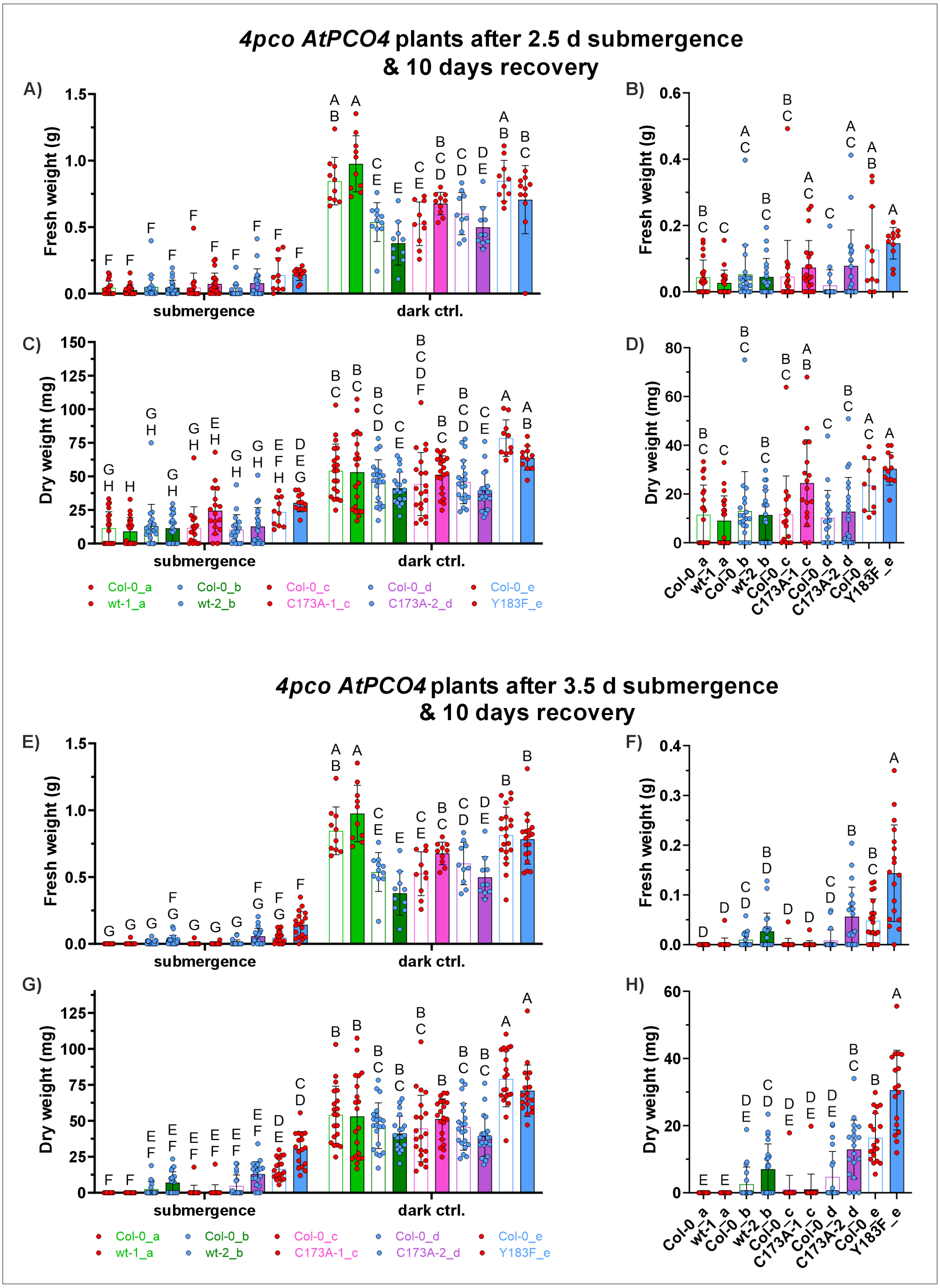


**Figure S19. Fresh and dry weight of *4pco AtPCO4 WT*, *C173A*, *Y183F* and Col-0 control plants after 2.5 and 3.5 d submergence.**

Fresh and dry weight of plants submerged for **(A)** – **(D)** 2.5 and **(E)** – **(H)** 3.5 days with **(A, C, E, G)** and without **(B, D, F, H)** dark control group after 10 days of recovery (lower case letters in legends indicate plants submerged in the same box). Dead plants were considered to weigh 0 g. The statistical significance ($P < 0.05$) was determined using two-way ANOVA followed by Tukey's multiple comparisons test (GraphPad Prism).

### Supplementary Tables

**Table S1. Primer sequences for *AtPCO4* mutagenesis.**
Primer names specify the encoded mutation and the name of a restriction enzyme which recognises an incorporated silent mutation.

| mutation | 5' – 3' |
|---|---|
| AtPCO4-1_Q134_A135insE_F | GATCCATCACAAGAAGCAAGACCTGCTAAACTGGTGAAGGATAC |
| AtPCO4-1_Q134_A135insE_R | GTTTAGCAGGTCTTGCTTCTTGTGATGGATCCTCTGGTTCG |
| **active site mutation** | |
| Y73R_XmiI_F | GCAATAAAGCGTCTACATTTGCATGAGTGTGACAGTTTC |
| Y73R_XmiI_R | GCAAATGTAGACGCTTTATTGCTGGGGGAGACTGATTAC |
| H98D_MamI_F | GATACCTCTTGATAACCATCCGGGCATGACCGTGCTAAG |
| H98D_MamI_R | GATGGTTATCAAGAGGTATCATAGAAGAAGGTGGCATACAGAAG |
| H100D_SmaI_F | CTTCATAACGACCCGGGCATGACCGTGCTAAG |
| H100D_SmaI_R | CATGCCCGGGTCGTTATGAAGAGGTATCATAG |
| V105G_F | CATGACCGGTCTAAGCAAGCTCGTTTATGGTTCAATG |
| V105G_R | CTTGCTTAGACCGGTCATGCCCGGATGGTTATGAAG |
| S107L_AflII_F | CATGACCGTGCTACTTAAGCTCGTTTATGGTTCAATGCATGTG |
| S107L_AflII_R | CATAAACGAGCTTAAGTAGCACGGTCATGCCCGGATGGTTATG |
| C173A_SphI_F | CAAAGCCATCACGCATGCGGCTATTCTTGACATCTTAGCTCCAC |
| C173A_SphI_R | CAAGAATAGCCGCATGCGTGATGGCTTTGAAACAGTGAATGTTG |
| I175F_NsbI_F | CATCACCCATTGCGCATTTCTTGACATCTTAGCTCCACCTTAC |
| I175F_NsbI_R | GTCAAGAAATGCGCAATGGGTGATGGCTTTGAAACAGTGAATG |
| D177E_BglII_F | CTATTCTTGAGATCTTAGCTCCACCTTACTCTTC |
| D177E_BglII_R | GCTAAGATCTCAAGAATAGCACAATGGGTGATG |
| Y183F_F[9] | ACATCTTAGCTCCACCTTTCTCTTCAGAGCATGATC |
| Y183F_R[9] | GATCATGCTCTGAAGAGAAAGGTGGAGCTAAGATGT |
| **reversion of spontaneous mutation in *AtPCO4-2* in pENTR_PCO4_AtPCO4-2** | |
| I148M_F | GATACTGAGATGACGGCTCAAAGCCCAGTAACAAC |
| I148M_R | CTTTGAGCCGTCATCTCAGTATCCTTCACCAGTTTAGC |

**Table S2**. **Primers for confirming complementation of *4pco* plants with *AtPCO4-2* WT and mutants.**

| primer | 5’ – 3’ | comment |
|---|---|---|
| attB1 | ACAAGTTTGTACAAAAAAGCAGGCT | Gateway® Technology, Invitrogen |
| attB2 | ACCACTTTGTACAAGAAAGCTGGGT | |
| PCO4_F | GCCTTACTTTGCTCAGAGGCTTTAC | |
| PCO4_R | CAAGTTCTAATGACAGGACCTCTGTACG | |
| pPCO4_F | CAGAATTCGCCCTTTTCACCGTTC | covering promotor region of |
| pPCO4_R | CACTAGTGATCTGAATCCAATTCAAAAG | *AtPCO4* |

**Table S3. Primer pairs for genotyping.**

| primer | 5' – 3' |
|---|---|
| insPCO5_F[10] | GCCCATTTAGGTAGCTGCAGTG |
| insPCO5_R[10] | AGCTTCCTGTTCGAGACCAA |
| LBb1 [10] | GCGTGGACCGCTTGCTGCAACT |
| promotorPCO4_F | ATGTGCGATTCATTTCCCAA |
| complPCO4_R | GGGCTTTGAGCCGTCATCTCAGTATCCTTC |
| PCO4_F | GCCTTACTTTGCTCAGAGGCTTTAC |
| PCO4_R | CAAGTTCTAATGACAGGACCTCTGTACG |

**Table S4. RT-qPCR primer sequences.**

| target | name | 5' – 3' | source |
|---|---|---|---|
| **Ubiquitin10 (At4g05320) (UBQ10)** | AT4G05320_F<br>AT4G05320_R | GGCCTTGTATAATCCCTGATGAATAAG<br>AAAGAGATAACAGGAACGGAAACATAGT | Licausi *et al*., 2011[11] |
| ***AtPCO4* (At2g42670)** | PCO4_KO_1_F<br>PCO4_KO_1_R | GAACGCAATGGAAGTAATCAGTCTC<br>TCATAGAAGAAGGTGGCATACAGAA | Dr Anna Dirr |
| | qpAtPCO4-2_F<br>qpAtPCO4-2_R | ACCAGAGGATCCATCACAAGAAGC<br>TACTGGGCTTTGAGCCGTCATC | Dr Anna Dirr |
| | sgPCO4_F<br>sgPCO4_R | CGAACCAGAGGATCCATCACAAGA<br>GCCGTCATCTCAGTATCCTTCACC | Prof Francesco Licausi |
| | PCO4_White_F<br>PCO4_White_R | CATGAGTGTGACAGTTTCTCTATAG<br>TGGTTCGGTCAGTTGAGGCTCTA | White *et al*., 2020[9] |
| ***AtPCO5* (At3g58670)** | AtPCO5_1_F<br>AtPCO5_1_R | CGACATCTTATCTCCTCCATACTCT<br>CATTCATCACTTCAATCTCACCAGG | Dr Anna Dirr |
| ***ADH1* (At1g77120)** | ADH_F<br>ADH_R | TATTCGATGCAAAGCTGCTGTG<br>CGAACTTCGTGTTTCTGCGGT | Licausi *et al*., 2011[11] |
| ***PDC1* (At4g33070)** | PDC1_F2<br>PDC1_R2 | GGACACCAAAATCGGATCGAT C<br>CTACTGAGGATTGGGAGGACG | Masson *et al*., 2019[10] |
| ***LBD41* (At3g02550)** | LBD41_F<br>LBD41_R | TGAAGCGCAAGCTAACGCA<br>ATCCCAGGACGAAGGTGATTG | Licausi *et al*., 2011[11] |
| ***SAD6* (At1g43800)** | SAD6_F<br>SAD6_R | TTGGCAACCCGCTTCTTTCTTACC<br>TTTCCCTCAGCTCACGAACCTG | Masson *et al*., 2019[10] |
| ***HB1* (At2g16060)** | HB1_F<br>HB1_R | ATGGAGAGTGAAGGAAAGATTGTG<br>TTAGTTGGAAAGATTCATTTCAGC | Gibbs *et al*., 2018[3] |
| ***SUS4* (At3g43190)** | SUS4_F<br>SUS4_R | AACGCAGAACGTGTAATAACG<br>CTCGGAGTGATGTTGAGTCC | Gibbs *et al*., 2018[3] |

**Table S5. Michaelis-Menten kinetics of *At*PCO4-2 enzymes using 0 – 1000 µM RAP.12$_{2–15}$.** Y183F values were determined from data for 0 – 2000 µM RAP.12$_{2–15}$ (Supplementary Figure S9C). ± indicates the SEM.

| | $V_{max}$ | | $K_M$ | | $k_{cat}$ | | $k_{cat}/K_M$ | |
|---|---|---|---|---|---|---|---|---|
| | µmol.mg$^{-1}$.min$^{-1}$ | | µM | | min$^{-1}$ | | min$^{-1}$.µM$^{-1}$ | |
| **WT** | **2.52** | **±0.07** | **100.30** | **±9.90** | **69.55** | **±1.97** | **0.69** | **±0.20** |
| **Y73R** | 0.04 | ±0.003 | 51.20 | ±12.28 | 1.12 | ±0.08 | 0.02 | ±0.006 |
| **V105G** | 0.29 | ±0.01 | 97.44 | ±15.54 | 7.94 | ±0.35 | 0.08 | ±0.02 |
| **S107L** | 0.49 | ±0.03 | 98.91 | ±18.60 | 13.61 | ±0.86 | 0.14 | ±0.05 |
| **C173A** | 2.51 | ±0.13 | 398.80 | ±46.85 | 69.20 | ±3.48 | 0.17 | ±0.07 |
| **I175F** | 1.81 | ±0.06 | 294.00 | ±23.32 | 50.01 | ±1.67 | 0.17 | ±0.07 |
| **D177E** | 0.13 | ±0.01 | 267.00 | ±55.23 | 3.53 | ±0.32 | 0.01 | ±0.006 |
| **Y183F** | 1.19 | ±0.09 | 1562.00 | ±206.40 | 32.90 | ±2.49 | 0.02 | ±0.01 |

**Table S6. Survival and recovery probability normalised to respective Col-0.**

| % compared to Col-0 | | | | | | |
|---|---|---|---|---|---|---|
| | | **wt-1** | **wt-2** | **C173A-1** | **C173A-2** | **Y183F** |
| **2.5 days** | **survival** | 50 | 83 | 145 | 300 | 125 |
| | **recovery** | 80 | 57 | 300 | 157 | 128 |
| **3.5 days** | **survival** | N/A | 275 | 100 | 233 | 118 |
| | **recovery** | N/A | 200 | 200 | 280 | 152 |

### Supplementary Discussion:

#### Additional discussion of the kinetic properties and predicted structural properties of *At*PCO4 variants

##### *Fe(II)-binding Residues: H98 and H100*

Our results confirmed that the His-triad is essential for *At*PCO4 activity as replacing H98 or H100 with an Asp diminished *At*PCO4 activity, to trace levels, similar to reported results for *At*PCO4-1 for H164D[9] and *At*PCO5 for H165[12]. *In silico* docking models of *At*PCO4 H98D and H100D (Figure S3) predicted a distorted iron coordination for both variants which potentially caused the reduced iron binding capacity and the depleted catalytic activity of *At*PCO4. Previous density functional theory studies of the thiol dioxygenase CDO predicted that mutating His-triad residues to Asp in CDO could alter $O_2$ activation and its transfer onto the Cys-thiol,[13] which may be consistent with the distorted iron coordination seen in these *At*PCO4 variants, H98D and H100D.

##### *Substrate-binding Residues: Y73 and Y183*

Mutating the substrate entry site residue Y73R also diminished the activity of *At*PCO4 almost completely (Figure 1). However, this correlated with a high $K_d$ and thus was due to reduced substrate affinity which suggested that Y73 is involved in substrate binding. The respective Tyr in *Hs*ADO, Y87, was proposed to affect *Hs*ADO activity due to modifying the positioning of the D206-carboxylate (equivalent residue in *At*PCO4: D177) via an H-bond network.[14] In *Hs*ADO, mutations Y87A and Y87F did not impact the $K_d$ considerably[14], suggesting that mutation Y73R in *At*PCO4 decreased substrate affinity mostly due to the change to a positively charged / basic side chain. The other residue at the proposed substrate entry site, Y183, was mutated to a Phe as examined previously by White *et al.*, 2020 in *At*PCO4-1[9] and by Chen *et al.*, 2021 in *At*PCO5[12]. The mutation Y183F in *At*PCO4 showed a similar reduction in activity as the previous results from *At*PCO4-1 and *At*PCO5 which can now be assigned to the decreased substrate affinity of the Y183F variant to RAP2.12$_{2-15}$. These findings correlate with the recent study of *Hs*ADO in which the authors mutated the respective Tyr residue to an Ala and Phe.[14] Both variants, *Hs*ADO Y212A and Y212F, were less active due to reduced substrate affinity.[14] This Tyr residue in *Hs*ADO is reported to be involved in a H-bond network with a water molecule and the Nt-Ser-OH of a substrate analogue (that was based on a cyclic peptide with a Gly-Ser arm reaching into the active site, PDB: 9DXB). An *At*PCO2 model with Cys as substrate reported by Chen *et al*., 2021[12] proposed that the Tyr at the equivalent position as Y183 in *At*PCO4 forms a H-bond with the Nt-Cys-$NH_3^+$. Either prediction for Nt-Cys coordination in the active site suggests an Nt-Cys stabilising function of *At*PCO4 Y183 which would correlate with the decrease in substrate affinity *in vitro* when the hydroxy group of the Tyr is removed. It remains to be determined if the Nt-Cys-SH or -$NH_3^+$ is stabilised by *At*PCO4 Y183.

***Residues trans to the His-triad: Asp177, Val105 and Ser107***

Whichever group on the Nt-Cys that is not stabilised by *At*PCO4 Y183, is likely stabilised by D177 which has been proposed by both studies[12,14]. In the *At*PCO2-Cys model[12], the small molecule substrate Cys-SH is predicted to form an H-bond with the *At*PCO2 D177-carboxyl group (*At*PCO4 numbering), whereas in the substrate analogue structure of *Hs*ADO the substrate's Nt-amine group interacts with the equivalent Asp-carboxyl via an H-bond (PDB: 9DXB). In addition, overlaying the *At*PCO4 structure with the *Hs*ADO substrate analogue structure suggests that D177 is within H-bonding distance to two water molecules that could mimic the position of $O_2$ coming into the active site and an $O_2$ molecule being coordinated by the metal cofactor. These structural insights and predictions confirm the kinetic results that residue D177 is important for catalysis (Figure 2). Extending the residue (by one -$CH_2$ group) while keeping the charge state ablated enzymatic activity by reducing the catalytic rate. However, the mutation D177E can still facilitate substrate binding at an affinity similar to the WT enzyme (Figure 2H). These results were in line with data from *Hs*ADO variants of the equivalent Asp residue.[14] Mutating the Asp to an Asn, as reported previously for *At*PCO4,[9] would result in a weaker H-bond between the Asn-amide and Cys-thiol groups than Asp-carboxylate and Cys-thiol, potentially, further reducing catalytic activity which has been observed when comparing both mutations in *Hs*ADO[14]. Due to a weakened H-bond the Asp-amide would not be able to stabilise the H-atom of the Cys-thiol as strongly, wherefore the Cys-S substrate would not be readily available for an attack by $O_2$. Another hypothesis could be that the Asp residue is necessary for $O_2$ activation or for the stabilisation of reaction intermediates. This theory is favoured for *Hs*ADO because the equivalent Asp in *Hs*ADO, D206, did not appear to deprotonate the Nt-Cys-thiol or -amine.[14]

The other two mutations *trans* to the iron-coordinating His-triad, V105G and S107L, also impacted the catalytic rate and catalytic efficiency of *At*PCO4 (Figure 2). Variant S107L appeared to have a lower catalytic rate and catalytic efficiency due to its reduced iron binding capacity and, potentially, modified positioning of the co-substrate $O_2$, reaction intermediate and/or the *At*PCO4 D177-carboxylate. Residue S107 could interact with the D177-carboxylate via an H-bond pulling the side chain into the active site and potentially towards the $O_2$ co-substrate. *Hs*ADO showed reduced activity when the equivalent Asp side chain (D206 in *Hs*ADO, D177 in *At*PCO4) was pulled towards the substrate entry site (PDB: 9DXU), i.e., in the opposite direction of *At*PCO4 S107, due to a cyclic peptide inhibitor[14]. However, the complete catalytic mechanism might be slightly different in *Hs*ADO than in *At*PCO4 because in *At*PCO4 the Ser at position 107 is replaced by a Leu in *Hs*ADO while the residue spatially below is a Val in *At*PCO4 but a Gly in *Hs*ADO (Figure S1C). These residues, Ser and Val in *At*PCO4 and Leu and Gly in *Hs*ADO, might affect the catalytic mechanism in both NCO enzymes by optimising the position for molecular $O_2$ coming into the active site in each enzyme (water molecules in *Hs*ADO substrate analogue structure PDB: 9DXB) and of the Asp residue that coordinates the protein and $O_2$ substrate. Reducing the size of S107 in *At*PCO4 in a previous study, S107A, did not affect *At*PCO4 activity[9] suggesting that a smaller but not larger residue is allowed at this position. Reducing the size of residue V105 with variant V105G, only affected the $V_{max}$ and catalytic rate and thereby the catalytic efficiency of *At*PCO4 but did not affect peptide substrate binding affinity (Figure 2). This further supported the hypothesis that residue V105 might be involved in optimising the position of the $O_2$ substrate in the active site by pushing it towards residues S107 and D177 by limiting the space elsewhere.

***Residues trans to the substrate entry site: C173 and I175***

Mutating a residue *trans* to the substrate entry site, C173 to A173, appeared to affect the catalytic rate of *At*PCO4 once $O_2$ was coordinated in the active site (Figure 3). The other residue *trans* to the substrate entry site that was investigated, I175 mutated to F175, showed RAP2.12$_{2\text{-}15}$ kinetics similar to variant C173A (Figure 3). The results supported the hypothesis that residues C173 and I175 are located closely to $O_2$ and/or reaction intermediates, and thus can affect $O_2$ activation and/or oxygen transfer onto the Nt-Cys-thiol. Examination of equivalent residues in *Hs*ADO in the recently reported crystal structure with bound substrate analogue (PDB: 9DXB) would suggest that both residues are likely to be too far away from the reaction centre to be involved in substrate binding. Any effects of the mutations might therefore be due to impacts on the *At*PCO4 structure or constrictions on conformational changes necessary for efficient catalysis.

### References in Supplementary


1. Honorato, R. V. *et al.* Structural Biology in the Clouds: The WeNMR-EOSC Ecosystem. *Front. Mol. Biosci.* **8**, 729513 (2021). doi:10.3389/fmolb.2021.729513.
2. Weits, D. A. *et al.* An apical hypoxic niche sets the pace of shoot meristem activity. *Nature* **569**, 714–717 (2019). doi:10.1038/s41586-019-1203-6.
3. Gibbs, D. J. *et al.* Oxygen-dependent proteolysis regulates the stability of angiosperm polycomb repressive complex 2 subunit VERNALIZATION 2. *Nat. Commun.* **9**, 5438 (2018). doi:10.1038/s41467-018-07875-7.
4. Labandera, A. *et al.* The PRT6 N-degron pathway restricts VERNALIZATION 2 to endogenous hypoxic niches to modulate plant development. *New Phytol.* **229**, 126–139 (2021). doi:10.1111/nph.16477.
5. Gunawardana, D. M., Southern, D. A. & Flashman, E. Measuring plant cysteine oxidase interactions with substrates using intrinsic tryptophan fluorescence. *Sci. Rep.* **14**, 1–10 (2024). doi:10.1038/s41598-024-83508-y.
6. Miles, A. J., Ramalli, S. G. & Wallace, B. A. DichroWeb, a website for calculating protein secondary structure from circular dichroism spectroscopic data. *Protein Sci.* **31**, 37–46 (2022). doi:10.1002/pro.4153.
7. Van Zundert, G. C. P. *et al.* The HADDOCK2.2 Web Server: User-Friendly Integrative Modeling of Biomolecular Complexes. *J. Mol. Biol.* **428**, 720–725 (2016). doi:10.1016/j.jmb.2015.09.014.
8. Madeira, F. *et al.* Search and sequence analysis tools services from EMBL-EBI in 2022. *Nucleic Acids Res.* **50**, W276–W279 (2022). doi:10.1093/nar/gkac240.
9. White, M. D. *et al.* Structures of Arabidopsis thaliana oxygen-sensing plant cysteine oxidases 4 and 5 enable targeted manipulation of their activity. *Proc. Natl. Acad. Sci.* **117**, 23140–23147 (2020). doi:10.1073/pnas.2000206117.
10. Masson, N. *et al.* Conserved N-terminal cysteine dioxygenases transduce responses to hypoxia in animals and plants. *Science* **365**, 65–69 (2019). doi:10.1126/science.aaw0112.
11. Licausi, F. *et al.* Oxygen sensing in plants is mediated by an N-end rule pathway for protein destabilization. *Nature* **479**, 419–422 (2011). doi:10.1038/nature10536.
12. Chen, Z. *et al.* Molecular basis for cysteine oxidation by plant cysteine oxidases from *Arabidopsis thaliana*. *J. Struct. Biol.* **213**, 107663 (2021). doi:10.1016/j.jsb.2020.107663.
13. de Visser, S. P. & Straganz, G. D. Why Do Cysteine Dioxygenase Enzymes Contain a 3-His Ligand Motif Rather than a 2His/1Asp Motif Like Most Nonheme Dioxygenases? *J. Phys. Chem. A* **113**, 1835–1846 (2009). doi:10.1021/jp809700f.
14. Jiramongkol, Y. *et al.* An mRNA-display derived cyclic peptide scaffold reveals the substrate binding interactions of an N-terminal cysteine oxidase. *Nat. Commun.* **16**, 4761 (2025). doi:10.1038/s41467-025-59960-3.
